# Predicted Bacterial uRBSs Reveal Translational Coupling and Ribosome-Mediated RBS Occlusion as Gene-Controlling Mechanisms

**DOI:** 10.64898/2026.05.12.723736

**Authors:** Theresa Dietz, Julian M. Hahnfeld, Sophie Neumann, Yonca A. Reinsch, Tessa Wenz, Susanne Barth-Weber, Jochen Blom, Alexander Goesmann, Elena Evguenieva-Hackenberg

**Author notes:** correspondence: **Corresponding author**: Elena Evguenieva-Hackenberg. These authors contributed equally to the study (Theresa Dietz, Julian M. Hahnfeld).

## Abstract

Upstream open reading frames (uORFs) in the 5′ leader of bacterial mRNAs can modulate gene expression, yet genome-wide identification remains limited. We combined bioinformatic prediction of ribosome-binding sites (RBSs) – a Shine-Dalgarno sequence and a start codon – with experimental validation to uncover new uORFs in *Sinorhizobium meliloti* 2011. From totally 1106 predicted upstream RBSs (uRBSs), we first examined 15 candidates using eGFP reporters and integrating existing RNA-seq and Ribo-seq data. Translation was detected at 13 sites, with fluorescence intensity broadly correlating with predicted initiation rates. Two uRBSs correspond to gene start sites, thereby refining gene annotations. In nine cases, uRBS mutations affected downstream gene expression in reporter fusions. Among others, the data suggests that a Type I secretion system operon, the RNA chaperone gene *hfq*, and metabolic genes are regulated by uORFs. Four uORFs acted through translational coupling. We also identified uRBSs that were ribosome-occupied yet (nearly) silent in eGFP assays, and closely spaced to the downstream main RBS (mRBS). These uRBSs probably mediate ribosomal occlusion downregulating *lacR* and SM2011_RS36230. A re-screen of the prediction set revealed 335 close uRBS/mRBS pairs. Three of them were analyzed, supporting the proposed ribosomal occlusion mechanism for SM2011_RS03630 and SM2011_RS22110, while for *glnK* translational coupling to an uORF was suggested. These results indicate that uORFs are more widespread in bacteria than previously recognized and suggest that direct ribosomal occlusion of the mRBS is a novel mechanism for down-regulating protein synthesis.

## INTRODUCTION

Upstream open reading frames (uORFs) are located in mRNA leaders and can play crucial roles in regulating gene expression in both eukaryotes and bacteria (Dar and Sorek, 2017; Dever et al., 2020). While their prevalence upstream of eukaryotic genes is well-documented (Bottorff et al., 2022; Dever et al., 2023; Zhang et al., 2019), knowledge on the occurrence of uORFs in bacterial genomes is still limited, preventing their functional analysis.

In eukaryotes, recent ribosome profiling (Ribo-seq) studies, in which occupancy of mRNA by ribosomes was determined, revealed that the majority of the protein coding genes are preceded by one or more uORFs (Bottorff et al., 2022; Zhong et al., 2024). Functional studies on eukaryotic uORFs highlighted their roles in development, cell homeostasis, circadian rhythm, and stress responses (Medenbach et al., 2011; Sun et al., 2025; Wiese et al., 2004; Young and Wek, 2016). Mostly, they act as *cis* regulatory elements that modulate the translation of the main ORF (mORF) by influencing the ribosome scanning (and thus the translation initiation efficiency) or the mRNA stability. Furthermore, some eukaryotic uORFs encode functional small proteins (Zhang et al., 2019).

Examples of bacterial uORFs, which were detected during analyses of specific genes, have been known for over 40 years (Dersch et al., 2017; Gemmill et al., 1979; Zurawski et al., 1978). They also can affect translation and stability of mORF transcripts, although mostly the mechanistic details differ from those in eukaryotic cells (Ben-Zvi et al., 2019; Dar and Sorek, 2017; Park et al., 2017). In addition, in the prokaryotic cell, uORFs can regulate the progress of transcription into downstream genes (Gemmill et al., 1979; Turnbough et al., 2019; Zurawski et al., 1978). Depending on the growth conditions, unhindered translation of an uORF (encoding a so-called leader peptide) or transient stalling at specific uORF codons determines the formation of mutually exclusive RNA structures or the accessibility of protein-binding sites in the mRNA leader. This, in turn, regulates the transcriptional or translational attenuation, or affects the mRNA-half-life and thus the mORF expression level (Ben-Zvi et al., 2019; Dar and Sorek, 2017; Turnbough et al., 2019).

Another, less explored function of bacterial uORFs that overlap the 5’-region of the associated mORFs is translational coupling (Park et al., 2017; Weaver et al., 2019). Translational coupling of neighboring genes is common in polycistronic mRNA, where translation of an upstream ORF enhances that of a downstream ORF. This typically occurs when ORFs have overlapping start and stop codons, or when the intergenic distance is short (usually less than 50 nucleotides). Mechanistically, this was explained by sliding 70S ribosomes, scanning 30S ribosomal subunit after dissociation of the 70S ribosome, and/or melting of secondary structures around the ribosome binding site (RBS) of the downstream ORF, which facilitates *de novo* binding of the 30S subunit (recently reviewed by Brown and Wade, 2025).

Finally, uORFs could encode small functional proteins. Increasing evidence that small proteins can have important physiological roles (Burton et al., 2024; Storz et al., 2014) forced attempts to identify and annotate translated small ORFs (sORFs) in prokaryotes by Ribo-seq (Bryant et al., 2023; Froschauer et al., 2025; Gelsinger et al., 2020; Hadjeras et al., 2023a; Hadjeras et al., 2023b; Laczkovich et al., 2022; Tufail et al., 2024). However, Ribo-seq is still not routinely applicable to different prokaryotic species (Vazquez-Laslop et al., 2022). Currently, it is best optimized for *E. coli*, where hundreds translated sORFs, among them potential uORFs, were uncovered in the last years (Meydan et al., 2019; Mohammad et al., 2019; Stringer et al., 2021; Weaver et al., 2019). Though, codon resolution typical for Ribo-seq of eukaryotes is often not achieved in prokaryotic studies. Among others, these experimental challenges impede the genome-wide detection of uORFs in bacteria.

In this study, we took advantage of the Shine-Dalgarno (SD) sequence present in the RBS of many translated ORFs in bacteria (Chen and Wen, 2022; Shine and Dalgarno, 1975; Wen et al., 2021) and predicted upstream RBSs (uRBSs) in *Sinorhizobium meliloti* 2011. We predicted 1123 uRBSs upstream of totally 1106 genes, studied 18 of them using reporter *egfp* fusions, and integrated existing RNA-seq and Ribo-seq data in the analysis (Hadjeras et al., 2023a; Kothe et al., 2025; Sallet et al., 2011). The results revealed new uRBSs and uORFs that increase or decrease the mORF expression. Interestingly, we uncovered uRBSs with strong peaks in the Ribo-seq, for which we could not detect translational products, but show that they downregulate the mORFs, most probably by mRBS occlusion. We suggest that these inhibiting uRBSs represent a novel mechanism for translational attenuation of gene expression by 70S ribosomes.

## MATERIAL and METHODS

### *In silico* detection of uRBSs and uORFs in *S. meliloti*

To identify novel uORF candidates in the *Sinorhizobium meliloti* 2011 genome, an *in silico* approach was developed to detect uORFs possessing own Shine-Dalgarno sequences (Fig. S1). These can form ribosomal binding sites with potentially high translation initiation rates (TIRs). The approach was implemented using Python (v3.12.9). Complete, unambiguous, and unfragmented coding sequences (CDSs) of the *S. meliloti* 2011 annotation (GCF_000346065.1-RS_2024_03_28) were extracted using gb-io.py (v0.3.4) (Larralde, 2025). For all extracted CDS the upstream region (−100 to +10 bp) was examined and uORFs as well as free start codons (uRBS) beginning with ATG, GTG, TTG, ATA, ATC, ATT, or CTG were kept for further evaluation. TIRs for the extracted uORFs were predicted using the OSTIR package (v1.1.2) (Roots et al., 2021) configured with the specific anti-Shine-Dalgarno (aSD) sequence of *S. meliloti*. All data were stored and processed using Polars (v1.24.0) (Vink et al., 2025) data frames. uORFs overlapping with upstream CDS were excluded from further analysis. The absolute delta of the predicted TIR for each uORF and its associated CDS was calculated (ΔTIR). Afterwards, Shine-Dalgarno sequence motifs with a length of 4 to 6 bp were extracted using a custom approach based on the aSD sequence of *S. meliloti* and the distance between the OSTIR predicted uRBS and the start codon of an uORF. In order to reduce the number of uORFs for each CDS, the uORF with the highest predicted TIR was kept with an additional lower threshold based on the first quantile of the predicted TIRs of all annotated CDS. This resulted in 912 uRBS candidates (Table S1). After manual inspection, eleven candidates were selected from this list of likely expressed uORF candidates for experimental analysis: Six candidates with a high TIR and delta (uSD1-6) and three candidates with a low TIR and delta (uSD7-11). The selected candidates also had to be in a 5’-UTR with a transcription start site (TSS) (Kothe et al., 2025; Sallet et al., 2011; the latter corresponds to the *S. meliloti* 2011 GenBank 2014 annotation harboring annotated 5’-UTRs).

To select additional candidates with a more conservative filter, all uORFs with a spacer sequence length between 4 and 10 bp and strongly conserved Shine-Dalgarno sequence motif consisting of “AGGAG” or “GGAGG” were retrieved, yielding 211 candidates with strongly conserved Shine-Dalgarno sequences (Table S2). From these, four additional candidates (uSD12-16) were selected for experimental analysis.

Furthermore, the predicted uORF positions were compared to publicly available Ribo-seq data and REPARATION and DeepRibo predictions (Ndah et al., 2017, Clauwaert et al., 2019) from the study of Hadjeras et al. (2023a) using polars.

The script for the uORF candidate prediction is available at https://github.com/jhahnfeld/ShineDal. Table S3 provides all software packages and tools used, along with their respective versions.

### Bacterial strains and their cultivation

*Sinorhizobium meliloti* 2011 was cultivated in rich TY medium (Beringer, 1974) or in GMS minimal medium (MM) (Schlüter et al., 2010). Routinely, 30 ml-cultures were shaken in 50 ml-Erlenmeyer flasks at 30°C and 140 r.p.m. Cells were harvested at OD_600nm_ of 0.5. Media were supplemented with 250 μg/ml streptomycin, and, when pRS2-derivatives were used, 10 µg/ml gentamicin. *Escherichia coli* DH5α and S17-1 strains were cultivated aerobically in LB broth with appropriate antibiotics (ampicillin 200 µg/ml; gentamicin 10 µg/ml) at 37°C and used for standard cloning or conjugation of shuttle plasmids into *S. meliloti* (Simon et al., 1983), respectively.

### Reporter plasmids and their construction

As a read-out for ORF expression, translational *egfp* fusions on plasmids were used. The *egfp* sequence was adapted to the *S. meliloti* codon usage (McIntosh et al., 2008). All used plasmids are listed in Table S4. The reporter plasmids contained short inserts in the shuttle plasmid pRS2, a gentamicin resistance mediating plasmid containing a pBBR backbone, a strong *metA* promoter (P*_metA_*), an *‘egfp* sequence lacking the first two *egfp* codons, and a strong heterologous terminator (Scheuer et al., 2022). Between P*_metA_* and *‘egfp*, XbaI and BamHI restriction sites are present, but a SD sequence is lacking. Using XbaI and BamHI, an mRNA leader and the first codons of an ORF can be cloned in frame with *egfp*, allowing to test the effect of the mRNA leader on translation of the ORF of interest. Inserts were prepared either by PCR or by annealing of two complementary oligonucleotides. Mutant derivatives were prepared either by cloning sequences containing the desired mutations, or by site-directed mutagenesis. Used oligonucleotides were purchased from Microsynth (Balgach, Switzerland) and are listed in Table S5. Enzymes were purchased from Thermo Fischer Scientific. Inserts were first cloned in plasmid pJET1.2/blunt (Thermo Fischer Scientific) and then subcloned into pRS2.

### Fluorescence measurements

*S. meliloti* strains, each containing one of the *egfp* reporter plasmids, were grown in TY or MM to OD_600 nm_ of 0.5, along with the empty vector control (EVC) strain harboring pRS2. Samples (150 µl aliquots) of the cultures were transferred to a 96-well black flatbottom microtiter plate. Fluorescence (extinction at 488 nm and emission at 522 nm) was measured using a Tecan Infinite M200 reader. Fluorescence values were normalized to the measured OD_600 nm_ and the EVC autofluorescence. At least three independent cultures in technical triplicates were used for the measurements. For each reporter plasmid, mean fluorescence and standard deviation were calculated. If fluorescence mediated by a reporter plasmid was not significantly higher than the EVC autofluorescence, the strain was considered non-fluorescent and its fluorescence was set to 1 (which is close to 0, since all statistically significant fluorescence values were above 200 arbitrary units).

### RT-qPCR

Steady-state amount of mRNA was quantified using Luna Universal One-Step RT-qPCR Kit (NEB). The used oligonucleotides are given in Table S5. RNA was purified with RNeasy (Qiagen), residual DNA was removed by treatment of 5 µg RNA with 0.5 µl TURBO-DNase (Invitrogen), and RT-qPCR was performed in a spectrofluorometric thermal cycler (Bio-Rad). BioRad CFX Manager 3.0 was used to determine the quantification cycle (Cq; the cycle, at which the amplification maximum of the curvature was reached) and the primer pair efficiencies in each PCR reaction. The levels were calculated in relation to the reference mRNA *rpoB*. Four independent experiments with technical duplicates were performed.

### Data visualization and statistical analysis

For graph preparation, Origin or GraphPad Prism were used. Statistical significance was determined using a *t*-test (Student, 1908), calculated by Microsoft Excel (Microsoft office LTSC Standard 2021). Pearson and Spearman correlation coefficients, corresponding p-values, and the confidence interval for the Pearson correlation were calculated using GraphPad Prism 8.4.3. Schematic drawings were prepared in PowerPoint. Existing Ribo-seq data (Hadjeras et al., 2023a) and RNA-seq data for TSS determination (Kothe et al., 2025) were visualized using the integrated genome browser (IGB).

## RESULTS

### Predicted uRBSs and uORFs

We first predicted 912 uRBSs consisting of a SD sequence and a start codon, which are located up to 100 nt upstream of annotated protein-coding genes, but not in coding sequences on the same DNA strand (Table S1). Due to this definition, many of the uRBS candidates are located in mRNA leaders, but candidates located in intergenic regions of polycistronic transcripts were also predicted, in line with previous observations (Choi et al., 2021). Each of the uRBS candidates in Table S1 has a calculated translation initiation rate (TIR) that is higher than the TIR of the associated, downstream annotated gene, which is designated a major ORF (mORF). Among the predicted uRBSs was a recently identified one (uRBS7 in Table 1), located in a riboswitch upstream of SM2011_RS30600 (*metA*; Corbino et al., 2005; Scheuer et al., 2022), supporting the reliability of our prediction. Additional prediction was conducted to highlight uRBS candidates containing the SD sequences AGGAG or GGAGG. This led to 211 predicted uRBSs located upstream of totally 194 mORFs (Table S2). Upstream of 8 mORFs, more than one putative start codon was assigned to the strong upstream SD (uSD). For the uRBSs containing AGGAG or GGAGG, we did not require higher TIR compared to the mORF. Only 50 mORFs are present in both Table S1 and Table S2. In sum, we predicted 1123 uRBSs upstream of totally 1106 genes.

**Table 1.** Experimentally analyzed uRBSs (uORFs) and corresponding mORFs. TIR: translation initiation rate; ΔTIR: TIR_uRBS_-TIR_mRBS_; Chr.: Chromosome; TR: transcriptional repressor; N.a.: Not applicable; the start codon in the uRBS is in frame with the mORF start codon and there is no stop codon between them. ^1)^ Listed in Table S2 with strong uSD; ^2)^ Longer isoform of a hypothetical protein (HP) encoded by the 2nd ORF of a bicistronic operon. ^3)^ Result may reflect a role of 4G-sequences rather than of an uRBS. ^4)^ Previously identified uRBS. The leader contains translation-activating uSD and a transcriptional terminator between uRBS and mRBS (Corbino et al., 2005; Scheuer et al., 2022).

| uRBS | Replicon | uRBS TIR | ΔTIR | uORF/mORF overlap | uORF start | mORF start | mORF identifier | mORF description | uRBS (uORF) function |
| --- | --- | --- | --- | --- | --- | --- | --- | --- | --- |
| 1 | Chr. | 4960.55 | 4419.68 | ATGA | ATG | ATG | SM2011_RS24730 | Hemolysin homolog | Translational coupling |
| 2 | pSymB | 3041.24 | 2982.36 | > 13 nt | GTG | ATG | SM2011_RS14295 | *araC* TR | Not clear |
| 3<sup>1)</sup> | Chr. | 2512.99 | 2407.37 | No overlap | ATG | GTG | SM2011_RS18345 | Oxidoreductase | Repressing (in TY) |
| 4<sup>1)</sup> | Chr. | 11682.65 | 11463.65 | No overlap | TTG | ATG | SM2011_RS25645 | Aldehyde dehydrogenase | Repressing (in MM)<sup>3)</sup> |
| 5 | Chr. | 10175.89 | 10017.04 | n.a. | ATG | ATG | SM2011_RS30165 | Phage TR | TIS of mORF |
| 6 | pSymB | 5115.62 | 5112.71 | n.a. | GTG | ATG | SM2011_RS08555 | TR | TIS of mORF |
| 7 | Chr. | 51.80 | 3.91 | n.a. | ATG | ATG | SM2011_RS30600 | *metA* | Activating uRBS<sup>4)</sup> |
| 8 | Chr. | 46.78 | 14.68 | No overlap | GTG | ATG | SM2011_RS21790 | *hfq* | Activating uORF |
| 9 | Chr. | 61.90 | 26.28 | 13 nt | TTG | ATG | SM2011_RS14345 | FxsA family protein | Translational coupling |
| 10 | pSymB | 43.18 | 29.23 | 375 nt | ATG | ATG | SM2011_RS14300 | *lacR* TR | Repressing uRBS |
| 11 | Chr. | 61.90 | 38.63 | ATGA | TTG | ATG | SM2011_RS25340 | *mcpW* | Not clear |
| 12<sup>1)</sup> | Chr. | 289.33 | 220.68 | Full overlap<sup>2)</sup> | ATG | ATG | SM2011_RS36230 | HP | Repressing uRBS |
| 13<sup>1)</sup> | pSymA | 382.66 | 380.38 | No overlap | ATG | ATG | SM2011_RS02580 | Mechanosensitive channel protein | Not clear |
| 14<sup>1)</sup> | pSymB | 1156.60 | 1156.6 | ATGA | ATG | ATG | SM2011_RS12745 | L-arabinose isomerase | Translational coupling |
| 15<sup>1)</sup> | Chr. | 277.98 | 275.19 | 68 nt | ATG | ATG | SM2011_RS15670 | GIY-YIG nuclease | Not clear |
| 16<sup>1)</sup> | pSymA | 326.72 | 310.48 | ATGA | ATG | ATG | SM2011_RS00160 | Gluconolactonase | Translational coupling |

Each uRBS candidate corresponds to a potential translation initiation site (TIS) of an uORF. Sequences of uORFs that do not overlap with the associated mORF (non-overlapping uORFs), or overlap the mORF by maximally 13 nt (Fig. 1A) are given in Tables S1 and S2. Putative larger overlaps were not determined, and corresponding uRBSs candidates are listed in Tables S1 and S2 only with the start codons. They represent two groups, out of frame or in frame with the mORF. Those of them, which are not in frame, represent uORFs overlapping their mORFs by > 13 nt. The second, in frame group, could represent misannotated or alternative TISs (Fig. 1A). The distribution of uRBSs belonging to the above categories is shown in Fig. 1B. Finally, Fig. 1C shows that most predicted uRBSs have ATG or TTG as a start codon, while GTG is less represented. There were no uRBSs with other alternative start codons among the predicted candidates.

**Figure 1.**
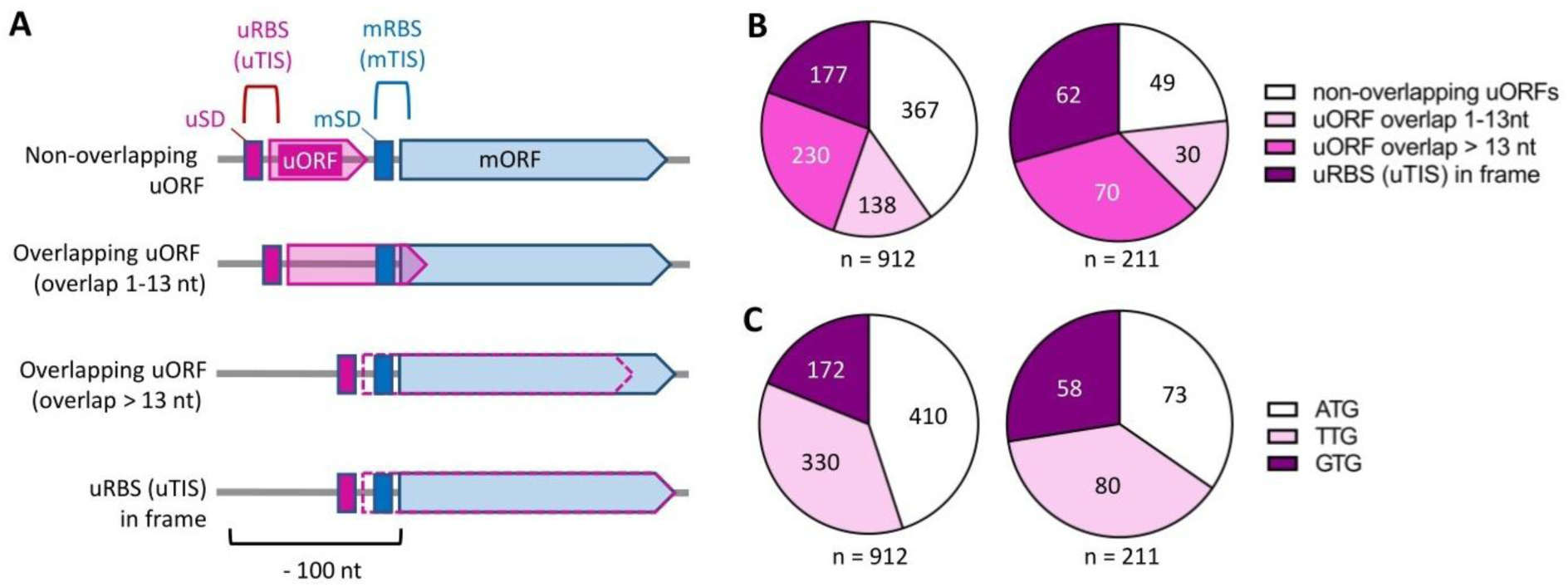
Overview of predicted upstream ribosome binding sites (uRBSs) and upstream open reading frames (uORFs) in *Sinorhizobium meliloti* 2011. **A)** Scheme of positions of uRBSs and uORFs in relation to the associated main ORFs (mORFs). The uRBS is defined as a Shine-Dalgarno sequence (SD) and a start codon. Each predicted uRBS corresponds to a potential upstream translation initiation site (uTIS). Depicted are four categories of uRBSs: 1) belonging to uORFs that do not overlap the mORFs; 2) belonging to uORFs that overlap the mORFs by maximally 13 nt; 3) belonging to uORFs that overlap the mORFs by more than 13 nt; 4) having a start codon in frame with the start codon of the mORF, without a stop codon between them. The latter are potential misannotated or alternative mORF start codons. mRBS, mTIS: the RBS and TIS of a mORF. **B)** Proportion of the four uRBS categories in Table S1 (n=912) and Table S2 (n=211). **C)** Proportion of the start codons ATG, TTG and GTG among the predicted uRBSs listed in Table S1 (n=912) and Table S2 (n=211). Table S1 lists uRBS candidates with a calculated TIR higher than the TIR of the associated mORF. Table S2 lists uRBS candidates containing the SD sequences AGGAG or GGAGG.

### Correlation between predicted TIR and ORF translatability in *Sinorhizobium meliloti*

We selected 15 new uRBS candidates for experimental analysis. Together with the known uRBS upstream of *metA,* they are listed in Table 1. To avoid uRBS candidates located outside of mORF transcripts, previously mapped transcription start sites (TSSs) (Kothe et al., 2025; Sallet et al., 2011) were considered. Further, the uRBS TIR and the difference to the TIR of the associated mRBS (ΔTIR) were considered and candidates with high (> 1000), moderate, and low (< 100) values were chosen. The chosen candidates (Table 1) represent the different uRBS (uORFs) categories shown in Fig. 1B and 1C. None of the selected uORFs have known homologs in either sORFdb (Hahnfeld et al., 2025), a specialized, taxonomy-independent small protein and sORF database for bacteria, or in GMSC (Duan et al., 2024), a catalog of small proteins from the global bacterial microbiome.

To test the translatability from the 15 new uRBSs and the RBSs of the corresponding mORFs (mRBSs), translational uORF-*egfp* and mORF-*egfp* fusions were cloned on plasmid pRS2 under the control of a strong heterologous promoter. In the constructs, each predicted start codon of an uORF or the annotated start codon of an mORF is preceded by a 21 nt sequence harboring the respective SD (Fig. 2A). An exception are uORF10-*egfp* and uORF12-*egfp*, for which the native TSSs were considered (the mRNA leader of uORF10-*egfp* was 28 nt and that of uORF12-*egfp* only 10 nt in length). *S. meliloti* 2011 strains containing the reporter plasmids were grown in rich medium (TY) and in minimal medium (MM), and fluorescence was measured.

**Figure 2.**
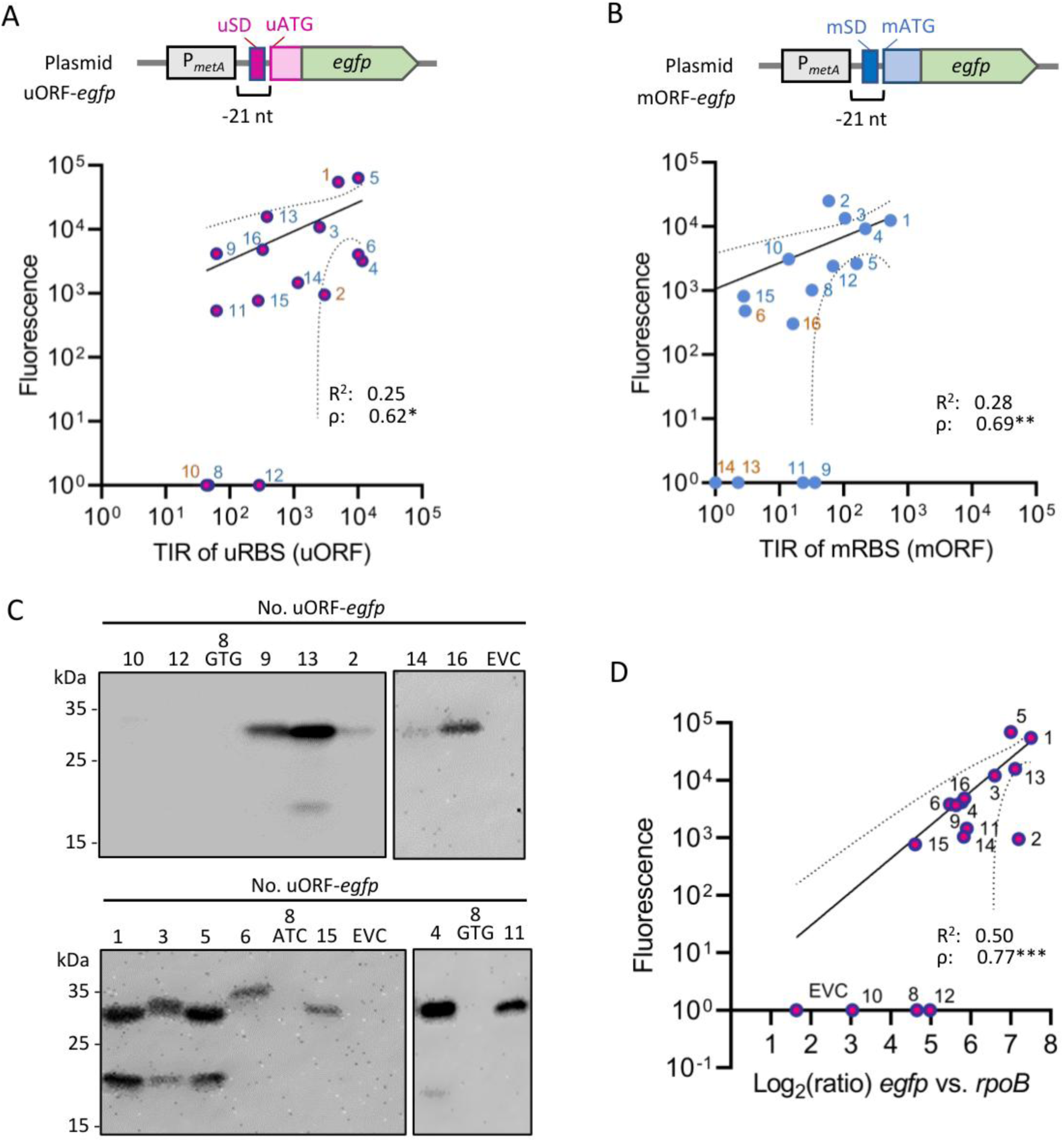
Correlation between predicted translation initiation rate (TIR) and fluorescence as a read-out of ORF translatability in *Sinorhizobium meliloti*. **A)** Analysis of 15 predicted uRBSs that represent putative uORF translation starts. Top: Scheme of the translational uORF-*egfp* fusions used for fluorescence (F) measurement. Bottom: Mean F values of bacterial cultures in rich TY medium, plotted against the predicted translation initiation rate (TIR) for each uRBS (uORF). For uRBS numbering, see Table 1. **B)** Analysis of the corresponding annotated mORFs. Top: Scheme of the used translational mORF-*egfp* fusions. Bottom: Mean F values of bacterial cultures grown in TY medium, plotted against the predicted TIR of each mORF. Data for bacterial cultures grown in minimal medium (MM) are shown in Fig. S2. For other descriptions see Fig. 1. Numbers in brown indicate constructs, for which the F-values in TY and MM show at least 2-fold difference. **C**) Western blot analysis of the uORF-*egfp* fusions in TY cultures using monoclonal GFP-directed antibodies. 8GTG corresponds to uRBS8. For 8ATC, see Fig. 6 below. **D)** Mean F values of the uORF-*egfp* fusions plotted against the relative amount of reporter mRNA, which was measured by RT-qPCR with *egfp*-directed primers. Internal control was the *rpoB* mRNA. The data were obtained from cultures grown in TY. In all graphs, the mean F values represent data from three independent biological experiments, each measured in triplicates. In case the mean F of a reporter construct was not significantly different from that of the empty vector control, F was set to 1. For RT-qPCR, four biological experiments were performed in technical duplicates. Results of Pearson and Spearman correlation analyses are shown. Coefficient of determination R^2^ and Spearman‘s Rho coefficient are given, and corresponding p-values are indicated: *** p ≤ 0.001, ** p ≤ 0.01, * p ≤ 0.05. Pearson regression line is shown and the confidence interval is indicated with dashed lines.

To address the correlation between TIR and fluorescence, mean fluorescence values were plotted against the predicted TIR values. Moderate linear correlation was detected by Pearson analysis, and Spearman analysis indicated strong relationship between TIR and fluorescence (Fig. 2A and 2B; Fig. S2A and S2B; the uRBS numbering is given in Table 1). Further, the data suggested condition-dependent expression of uORF10 and mORFs number 6, 13, 14, and 16 (showing significant fluorescence only in TY or MM), and for uORF1 and uORF2 (showing > 2-fold difference in fluorescence between TY and MM; compare Fig. 2 to Fig. S2). Additionally, for uORFs 8 and 12, as well as for mORFs 9 and 11, no fluorescence was detected under both conditions (compare Fig. 2 to S2). We note that although for mORFs 6, 9, 11, 13, 14, and 16 no fluorescence was detected under at least one condition, the corresponding uORFs showed fluorescence in TY and MM (Fig. 2 and Fig. S2), suggesting that these mORFs could be activated by the corresponding translated uORFs.

The lack of strong linear correlation between TIR and fluorescence could be explained by several factors: 1) The GC-rich genome of *S. meliloti* is suboptimal for the used TIR prediction model of OSTIR, as it was primarily developed for *Escherichia coli*; 2) inactive aggregates of overproduced eGFP distort fluorescence results; 3) differences in protein and mRNA stability among the constructs affect the produced eGFP amount independently of the TIR. To address some of these factors, we conducted western blot (Fig. 2C) and RT-qPCR (Fig. 2D) analyses of TY cultures harboring the uORF-*egfp* constructs. Relative signal intensities (as visually estimated), or lack of signals were in line with the fluorescence data (compare Fig. 2C to Fig. 2A), suggesting that fluorescence reflects the eGFP amounts. Further, the levels of the individual uORF-*egfp* mRNAs, which are transcribed from the same strong promoter, correlated better with the fluorescence intensity than the predicted TIRs (compare Fig. 2D to Fig. 2A and 2B). However, the data do not allow to distinguish between inherent differences in mRNA stability among the constructs and differences due to ribosome binding and translation (Deana and Belasco, 2005).

Moreover, we used all annotated ORFs, for which Hadjeras et al. (2023a) detected translation by Ribo-seq, to predict their TIRs, but only a weak correlation was detected between Ribo-TE at the region covered by the initiating ribosome and predicted TIR (Pearson: R = 0.143, Spearman: Rho = 0.294). This could, at least in part, be explained by the fact that the TIR prediction by OSTIR did not consider regulation in the cell by RNA-binding proteins and regulatory RNAs.

Altogether, for 13 out of the 15 newly tested uRBSs, significant fluorescence of uORF-*egfp* was detected under at least one condition (Fig. 2A and Fig. S2A). This suggests that many of the predicted uRBSs probably drive uORF translation in *S. meliloti*.

### Predicted uRBS5 and uRBS6 harbor start codons of SM2011_RS30165 and SM2011_RS08555

The new candidates uRBS5 and uRBS6 (Table 1) have start codons that are in frame with the mORFs, without a stop codon in between, suggesting that they are alternative or major translational starts. They are located upstream of SM2011_RS30165 and SM2011_RS08555, respectively, both encoding transcriptional regulators. For both genes, fluorescence of the uORF-*egfp* fusions was much higher than that of the mORF-*egfp* fusions (Fig. 3A and 3B). These results are in line with the high TIR and ΔTIR values of uRBS5 and uRBS6.

**Figure 3.**
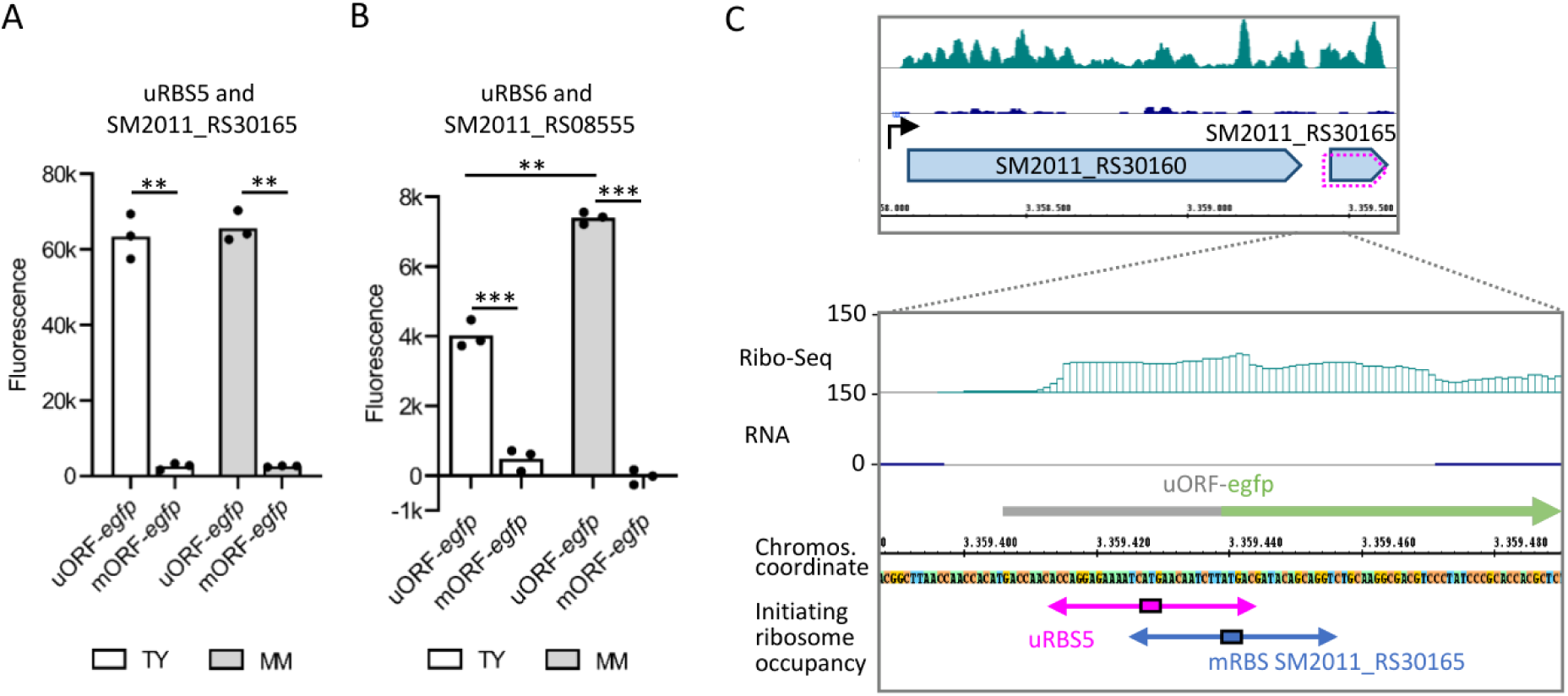
Predicted uRBS5 and uRBS6 are start codons of SM2011_RS30165 and SM2011_RS08555. **A)** Fluorescence mediated by the uORF-*egfp* fusion of the predicted uRBS5 and by the corresponding mORF-*egfp* (SM2011_RS30165-*egfp*) fusion in bacterial cultures grown in TY or MM. **B)** Fluorescence mediated by the uORF-*egfp* fusion of the predicted uRBS6 and by the corresponding mORF-*egfp* (SM2011_RS08555-*egfp*) fusion in bacterial cultures grown in TY or MM. All graphs show means and single data points of three independent experiments. Significance of difference determined by *t*-test: *** p ≤ 0.001, ** p ≤ 0.01, * p ≤ 0.05. **C)** Integrated genome browser (IGB) screenshots depicting reads from Ribo-seq and RNA-seq libraries. Shown is the SM2011_RS30160-SM2011_RS30165 operon and its transcriptional start site (TSS, marked by a bent arrow). A zoom-in on the intergenic region harboring uRBS5 and the mRBS of SM2011_RS30165 is also shown. Horizontal double-headed arrow: a 31 nucleotides (nt) region that is expected to be occupied by an initiating ribosome, which uses the start codon (small box) in the center of this region. In the Ribo-seq library, read enrichment shows ribosome occupancy at the ATG of uRBS5 as a start codon. The region cloned in the L-mORF-*egfp* construct is indicated by a horizontal gray arrow. IGB screenshots for uRBS6 and SM2011_RS08555 are shown in Fig. S3.

Available Ribo-seq data reflecting translation during growth in MM (Hadjeras et al., 2023a), support translational start of SM2011_RS30165 at uRBS5 (Fig. 3C). For SM2011_RS08555, translation was not detected by Ribo-seq, but the data suggest ribosome binding in the mRNA leader upstream of uRBS6, pointing to the existence of a non-overlapping uORF (Fig. S3). We note that this applies for a transcript with mRNA leader as annotated 2014 in the GenBank. Our recent TSS mapping in *S. meliloti* 2011 grown semiaerobically in TY (Kothe et al., 2025) revealed a leaderless SM2011_RS08555 transcript, which does not harbor uRBS6. Together, this suggests the existence of alternative TSSs yielding SM2011_RS08555 transcripts with different lengths, which are translated in protein isoforms. The larger isoform harbors an N-terminal MLDEL sequence, which provides negatively charged residues that could impact protein function. Fig. 3B suggests that the expression of the larger isoform is higher in MM than in TY.

To identify uRBSs that could represent misannotated or alternative translation initiation sites (TISs), we used the available Ribo-seq data, and REPARATION and DeepRibo predictions (Ndah et al., 2017, Clauwaert et al., 2019) from the study of Hadjeras et al., 2023a. This revealed 23 possibly missanotated candidates (Tables S6a and S6b). Additionally, Tables S6c and S6d show 15 Ribo-seq-supported uRBS that are likely TISs of separate uORFs.

### Overlapping uORFs that act by translational coupling

To address the roles of the 13 remaining, new uRBSs and their uORFs, we cloned L-mORF-*egfp* constructs harboring extended mRNA leaders (L) (Fig. 4A). Fluorescence yielded by these and the aforementioned mORF-*egfp* and uORF-*egfp* constructs (Fig. 4A) was measured. Then ratios R1 = F_uORF-egfp_/F_mORF-egfp_ and R2 = F_L-mORF-egfp_/F_mORF-egfp_ were calculated and plotted against each other (Fig. 4B and 4C). We hypothesized that uRBSs (uORFs) grouped in one of the quadrants (Q) I to IV in Fig. 4 have similar features and functions: QI refers to cases in which the uORF is highly translated relative to the mORF, and the mRNA leader enhances mORF expression, indicating a positive effect of the uORF. QII represents uORFs that are weakly translated, or not translated at all, yet are situated within an mRNA leader that enhances mORF translation. In this configuration, a conditional rise in uORF translation could influence mORF expression, or the binding of ribosomes to the uRBS may serve a regulatory role. QIII describes uORFs that are weakly translated, or not translated at all, and reside in an mRNA leader that suppresses mORF translation. The regulatory scenarios here are analogous to those in QII: a conditional uORF translation could alter mORF expression, or ribosome engagement at the uRBS might have regulatory function. Finally, QIV represents uORFs that are strongly translated relative to the downstream mORF and are located in mRNA leaders that repress mORF translation, indicating a negative regulatory effect by the uORF.

**Figure 4.**
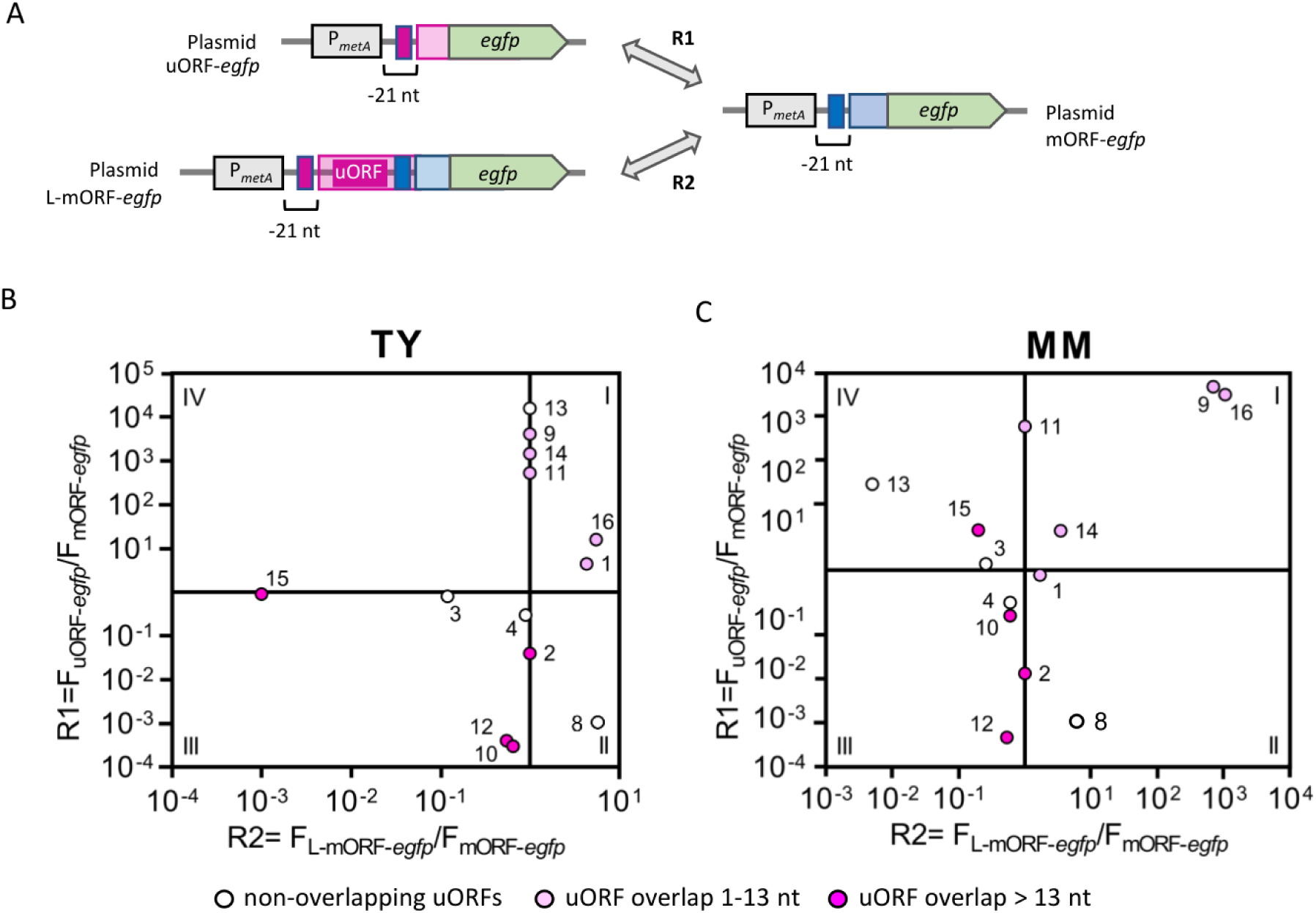
Most uRBS-containing mRNA leader affect mORF expression. **A)** Scheme of the used uORF-*egfp*, mORF-*egfp* and L-mORF-*egfp* constructs, the latter harboring mRNA leader (L) with uRBS. For other descriptions, see Fig. 1. **B)** and **C)** R1 = F_uORF-*egfp*_/F_mORF-*egfp*_ was plotted against R2 = F_L-mORF-*egfp*_/F_mORF-*egfp*_. Used growth medium is indicated. The points corresponding to each analyzed uRBS and its associated uORF and mORF are colored to mark the uORF category according to Fig. 1A. For uRBS/uORF/mORF numbering, see Table 1.

In QI, which contains uORFs with presumably positive effect on mORF expression, uORFs that overlap the mORFs with max. 13 nt were located: uORFs 1 and 16 in TY (Fig. 4B), and uORFs 9, 14, and 16 in MM (Fig. 4C; for numbering see Table 1). All except uORF9 show a start-to-stop (ATGA) overlap with the mORF, suggesting that they act by translational coupling (Brown and Wade, 2025). The uORF11 also has an ATGA overlap with *mcpW*, but was excluded from the further analysis, because the mORF-and L-mORF-*egfp* fusions did not yield significant fluorescence in TY nor MM, the uORF-*egfp* fluorescence was low, and Ribo-seq did not detect translation at this locus. For the remaining four overlapping uORFs, Ribo-seq suggested ribosome occupancy at the start codons (Fig. 5A; Fig. S4). To test whether translation of the uORFs impacts mORF translation, we studied the effect of start-to-stop mutations destroying the uORFs in the L-mORF-*egfp* constructs. As expected, the mutations decreased or abolished the mORF expression (Fig. 5B to 5E). Thus, we uncovered that the following genes are preceded by translationally coupled uORFs: SM2011_RS24730 (encodes a hemolysin homolog; a first gene in a T1SS operon preceded by uRBS1), SM2011_RS14345 (encodes FxsA family protein; preceded by uRBS9), SM2011_RS12745 (encodes L-arabinose isomerase; preceded by uRBS14), and SM2022_RS00160 (encodes SMP-30/gluconolactonase/LRE family protein; preceded by uRBS16). Further, we note that nearly 10 % of the predicted uORFs have ATGA overlaps with their mORFs (Table S7).

**Figure 5.**
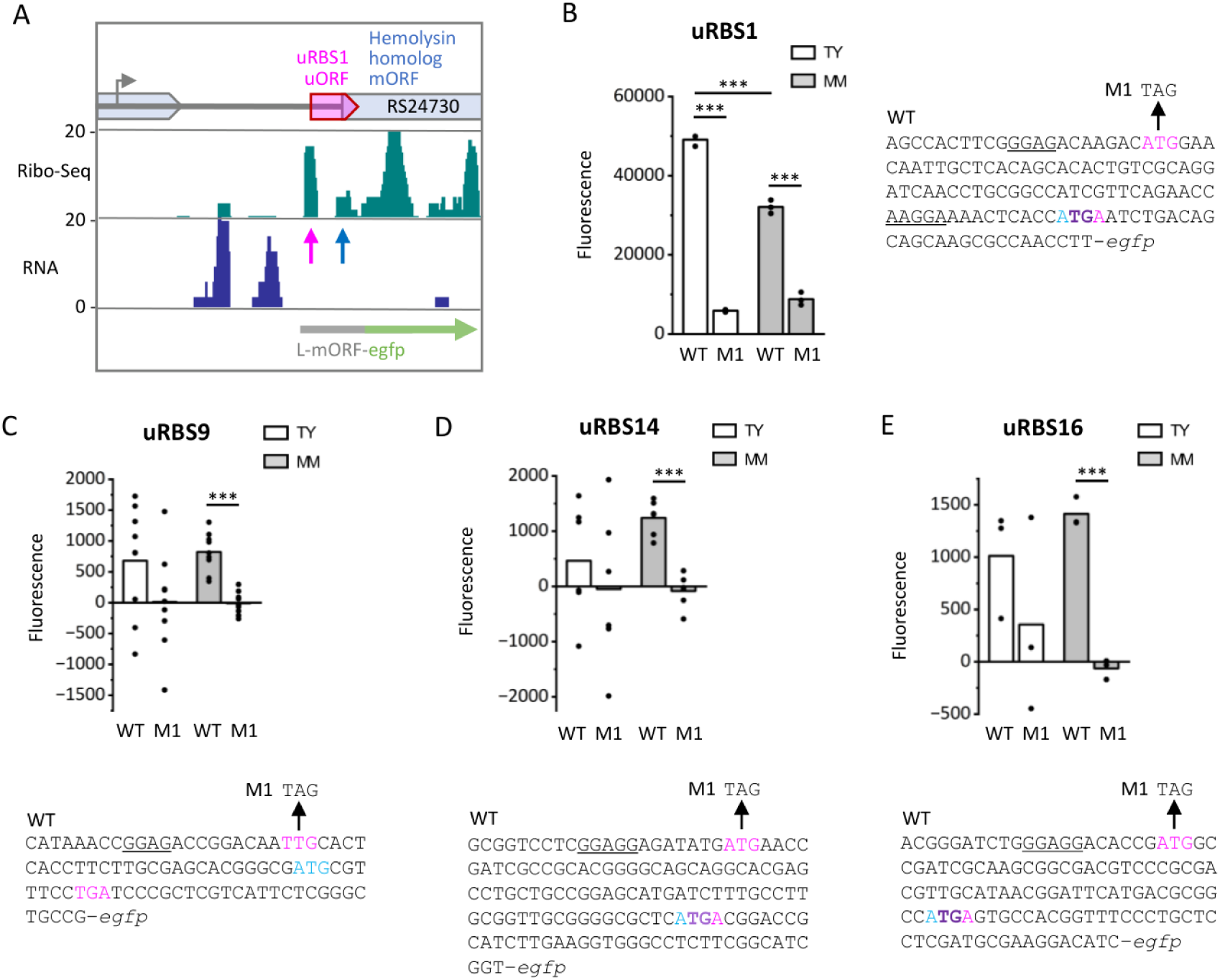
Upstream ORFs that increase mORF expression by translational coupling. **A)** Integrated genome browser (IGB) screenshots depicting reads from Ribo-seq and RNA-seq libraries for the region encompassing uRBS1 and the mORF SM2011_RS24730. Ribosome coverage peaks corresponding to the uRBS and the mRBS are indicated by pink and blue vertical arrows, respectively. Annotated genes are depicted in light blue and the overlapping uORF in pink. A transcription start site, located in an upstream annotated gene, is indicated by a bent gray arrow. IGB screenshots of uRBS9, 14, and 16 are shown in Fig. S4. **B) – E)** Fluorescence mediated by L-mORF-*egfp* fusions (WT) and their corresponding derivatives with start-to-stop mutations destroying the uORFs in the mRNA leader (M1). The uRBS numbers (see Table 1) and the used growth conditions are indicated. Cloned sequences are shown. Start and stop codons are in pink for the uORF and cyan for the mORF. Start-stop overlap in ATGA is indicated in purple. Shine-Dalgarno (SD) sequences that contain at least four nucleotides fully complementary to the anti-SD sequence of the *S. meliloti* 16S rRNA are underlined. All graphs show means and single data points of at least three independent experiments. Significance of difference determined by *t*-test: *** p ≤ 0.001, ** p ≤ 0.01, * p ≤ 0.05.

### uORFs that do not overlap the mORF: two uORFs in the *hfq*-activating mRNA leader

In Fig. 4 above, the non-overlapping uORFs numbered 3, 4, 8, and 13 are scattered in quadrants II to IV, suggesting diverse modes of action. For uORF3 and uORF4, both located in intergenic regions of polycistronic mRNA, Fig. 4 suggests a negative impact on mORF expression. For both of them, two L-mORF-*egfp* derivatives were cloned and analyzed, one with mutated uSD and a second one with destroyed uORF.

The 5-amino acid (aa) uORF3 is located upstream of SM2011_RS18345 (encoding a FAD-dependent oxidoreductase). An ATG/TAG mutation destroying the uORF in the L-mORF-*egfp* increased fluorescence in TY, but not in MM, suggesting a downregulating role of uORF3 under specific conditions. However, the uSD mutation had no significant effect on reporter expression, questioning the role of uORF3 (Fig. S5A and S5B).

The “startstop” uORF4 TTGTGA, located upstream of SM2011_RS25645 (encoding an aldehyde dehydrogenase family protein), has the highest predicted TIR and ΔTIR in Table S1. Replacement of TTGTGA by TTAAGA in L-mORF-*egfp* did not change reporter expression. However, the uSD-mutation led to a 2-fold increase of the reporter expression in MM only, suggesting a condition-specific, negative effect of 30S ribosomal subunit binding to the uSD (Fig. S5C and S5D). However, we note that this uSD is a part of a G-quadruplex (G4) region (Cueny et al., 2022). Thus, the uSD-mutation could interfere with a condition-specific function of the G4-motifs in regulation of SM2011_RS25645.

For the 5-aa uORF13, located in the mRNA leader of SM2011_RS02580 (encoding a mechanosensitive ion channel domain-containing protein), Fig. 4C suggests downregulating function in MM. Ribo-seq detected ribosome binding at the uRBS, but did not support mORF translation in MM (Fig. S5E). In both MM and TY, uORF-*egfp* showed strong fluorescence (see Fig. 2A and Fig. S2A above). However, no fluorescence was detected for L-mORF-*egfp*, and the ATG/TAG mutation destroying the uORF in L-mORF-*egfp* did not activate the reporter expression (Fig. S5F). Thus, the results strongly support uORF translation, but its impact on SM2011_RS02580 remains unclear.

Finally, the uORF8 located upstream of *hfq* was analyzed. According to Fig. 4, the uORF8-harboring mRNA leader increases *hfq* expression, despite no detectable uORF8 translation (see also Fig. 2A and Fig. S2A above). Close inspection of the Ribo-seq data revealed ribosome coverage that is centered around an alternative ATC start codon, upstream of the GTG start codon of uORF8 (Fig. 6A). The ATC-uORF (19 aa) overlaps the short GTG-uORF (4 aa). We tested the translation of both uORFs by uORF-*egfp* fusions and reproducibly obtained weak fluorescence only for ATC-uORF-*egfp* in MM (Fig. 6B). Since the ribosome at the ATC-RBS would occlude the GTG-RBS, ATC was mutated to TAC in the GTG-uORF-*egfp* construct, but the mutation had no significant effect (Fig. 6B). Next, to address the roles of both uORFs, we mutated their start codons in the L-mORF-*egfp* construct separately and together. Fig. 6C shows that only the GTG/GTC mutation destroying the originally predicted GTG-uORF has impact on *hfq* expression. This point mutation, alone or in combination with ATC/TAC, almost completely diminished the activating role of the mRNA leader on *hfq*-*egfp* expression (Fig. 6C). We conclude that the GTG-uORF could be necessary for efficient *hfq* expression, while the role of the ATC-uORF remains unclear.

**Figure 6.**
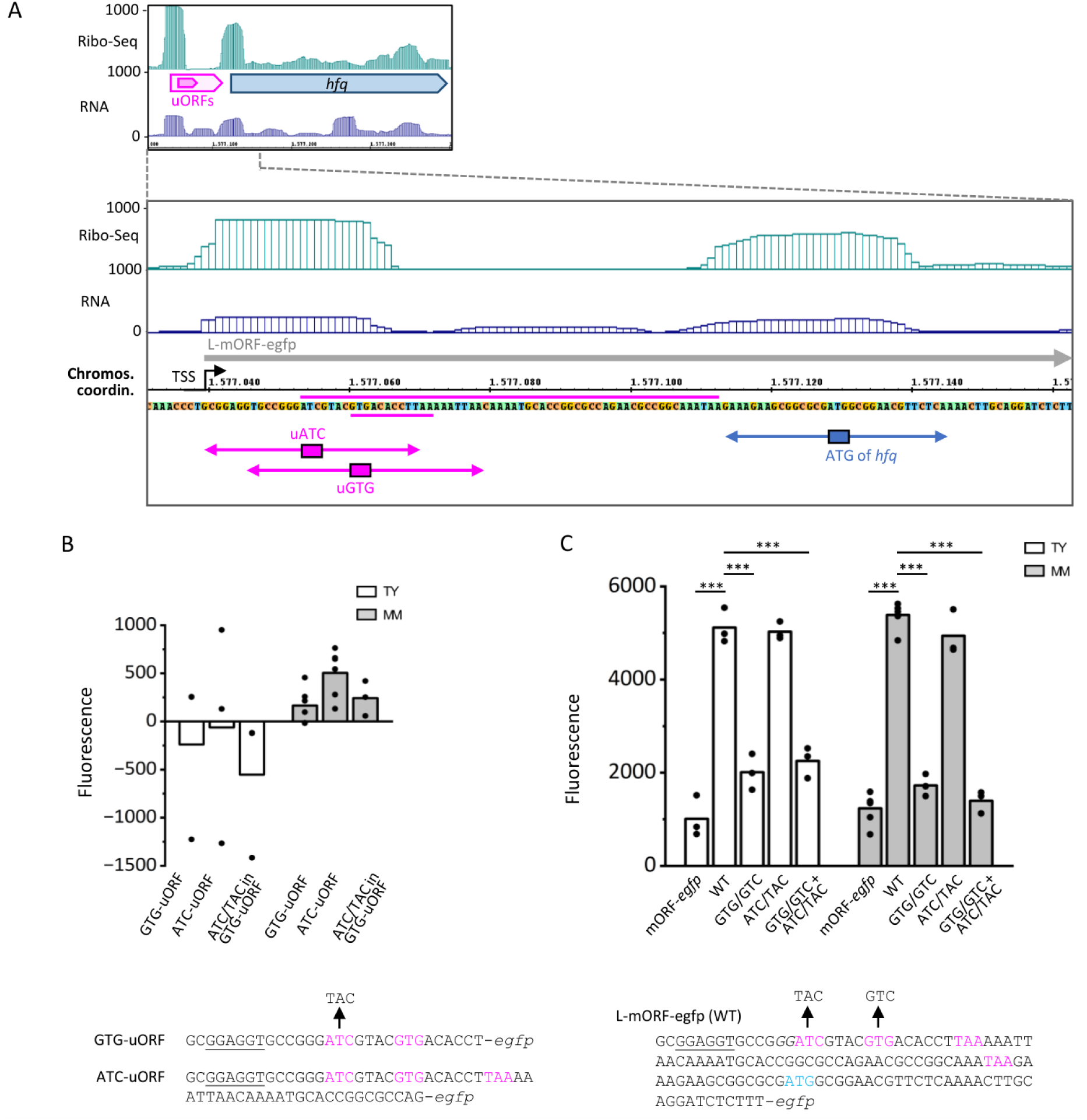
Point mutation in the mRNA leader reduces *hfq* expression, suggesting a role for uRBS8. **A)** Integrated genome browser screenshots depicting reads from Ribo-seq and RNA-seq libraries. The upper panel shows Ribo-seq peaks at the start codons of uORF(s) and of *hfq*. Two overlapping uORFs are depicted in pink and *hfq* in blue. The zoom-in indicates that the prominent peak in the mRNA leader aligns with ribosome occupancy at an ATC start codon, rather than at the GTG start codon predicted for uORF8. The regions corresponding to the two uORFs (the short GTG-uORF overlapped by the longer ATC-uORF) are marked by horizontal pink lines. For other descriptions, see Fig. 3. **B)** Fluorescence mediated by the indicated uORF-*egfp* fusions. **C)** Fluorescence mediated by mORF-*egfp*, L-mORF-*egfp* (WT), and by derivatives of the latter harboring mutations destroying the GTG-uORF (GTG/GTC), the ATC-uORF (ATC/TAC) or both mutations. Cloned sequences are shown. Start and stop codons are in pink for the uORFs and cyan for the mORF. Shine-Dalgarno (SD) sequences that contain at least four nucleotides fully complementary to the anti-SD sequence of the *S. meliloti* 16S rRNA are underlined. Growth conditions are indicated. All graphs show means and single data points of at least three independent experiments. Significance of difference determined by *t*-test: *** p ≤ 0.001, ** p ≤ 0.01, * p ≤ 0.05.

### Analysis of largely overlapping uORFs suggests that uRBS10 and uRBS12 act by mRBS occlusion

The uORFs numbered 2, 10, 12, and 15 overlap their corresponding mORFs by > 13 nt. Fig. 4 above suggests that, with the exception of uORF2, the other uORFs may negatively impact mORF expression.

A closer look at uORF2 shows that its GTG start codon (uGTG) and the mATG of the mORF SM2011_RS14295 (encodes AraC transcriptional regulator) share the same SD (Fig. S6A). A high TIR and ΔTIR were predicted for uRBS2, and thus we expected that a translational fusion in frame with uGTG will give much higher fluorescence than a fusion in frame with mATG. However, fluorescence measurements suggested weak translation from uGTG as the start codon and strong translation starting at the mATG (Fig. S6B). An mATG/TAG mutation in the uGTG-*egfp* fusion did not change fluorescence significantly (Fig. S6B), suggesting that there is no competition between the uGTG and mATG start codons, and that the lack of strong translation from the uGTG is an inherent feature of uRBS2. Next, we mutated the uGTG in the ATG-*egfp*-fusion. Since changes in the spacer between the SD and mATG are expected to influence mORF translation, we mutated the uGTG to either GTC, GTA, GTT, or AAA. Only the GTG/GTC mutation, which introduces an unfavorable nucleotide in the spacer (Kuo et al., 2020), had a significant negative effect (Fig. S6B). Altogether, the results argue against a role of uRBS2 and the associated 11-aa uORF, although translation from the GTG as a start codon was measurable and the data suggested differential uORF translation in TY and MM (Fig. 2A and S2A).

The uRBS15 is located in an alternative, longer transcript of the otherwise leaderless mORF SM2011_RS15670 (encodes a GIY-YIG nuclease family protein; Fig. S6C). The mORF-*egfp* showed fluorescence, but the L-mORF-*egfp* did not, suggesting repressive effect of the uORF-containing mRNA leader (Fig. S6D). However, mutation destroying uORF15 had no significant effect (Fig. S6D). Thus, it is not clear whether uRBS15 plays a role in SM2011_RS15670 expression.

The uORF10 largely overlaps with the mORF SM2011_RS14300 (*lacR*) (Fig. 7A). Fig. 2A and Fig. S2A above indicate that uORF10 is not translated in TY and is weakly translated in MM. We constructed several uORF10-*egfp* fusion variants addressing the translatability of uORF10 (Fig. 7SA). Their analysis in MM cultures suggested that the 3-nt spacer between uSD and uATG in uRBS10, and rare N-terminal codons limit uORF10-*egfp* translation (Fig. 7SB). In contrast to this data, Ribo-seq indicated that in MM, 70S ribosomes occupy uRBS10 with an intensity comparable to that at the mRBS (Fig. 7A). The uATG and the mATG of *lacR* are separated by only seven nucleotides (Fig. 7B) and this could lead to competition for and mutually exclusive occupancy by initiating ribosomes. To address this, we mutated the uATG to ACG, thereby inactivating uRBS10 in the L-mORF-*egfp* construct (M1 mutation in Fig. 7B; start-to-stop mutation was avoided because it would create an artificial mSD). The point mutation significantly increased fluorescence (Fig. 7C), in line with the expected lack of competition between uRBS and mRBS. According to the OSTIR prediction, the SD sequence AAGGA (underlined in the WT construct shown in Fig. 7B) is used by both the uATG and the mATG. Accordingly, deletion of this SD in the L-mORF-*egfp* construct (M2 mutation in Fig. 7B) resulted in lower fluorescence (Fig. 7C). Altogether, the results support inhibiting role of uRBS10.

**Figure 7.**
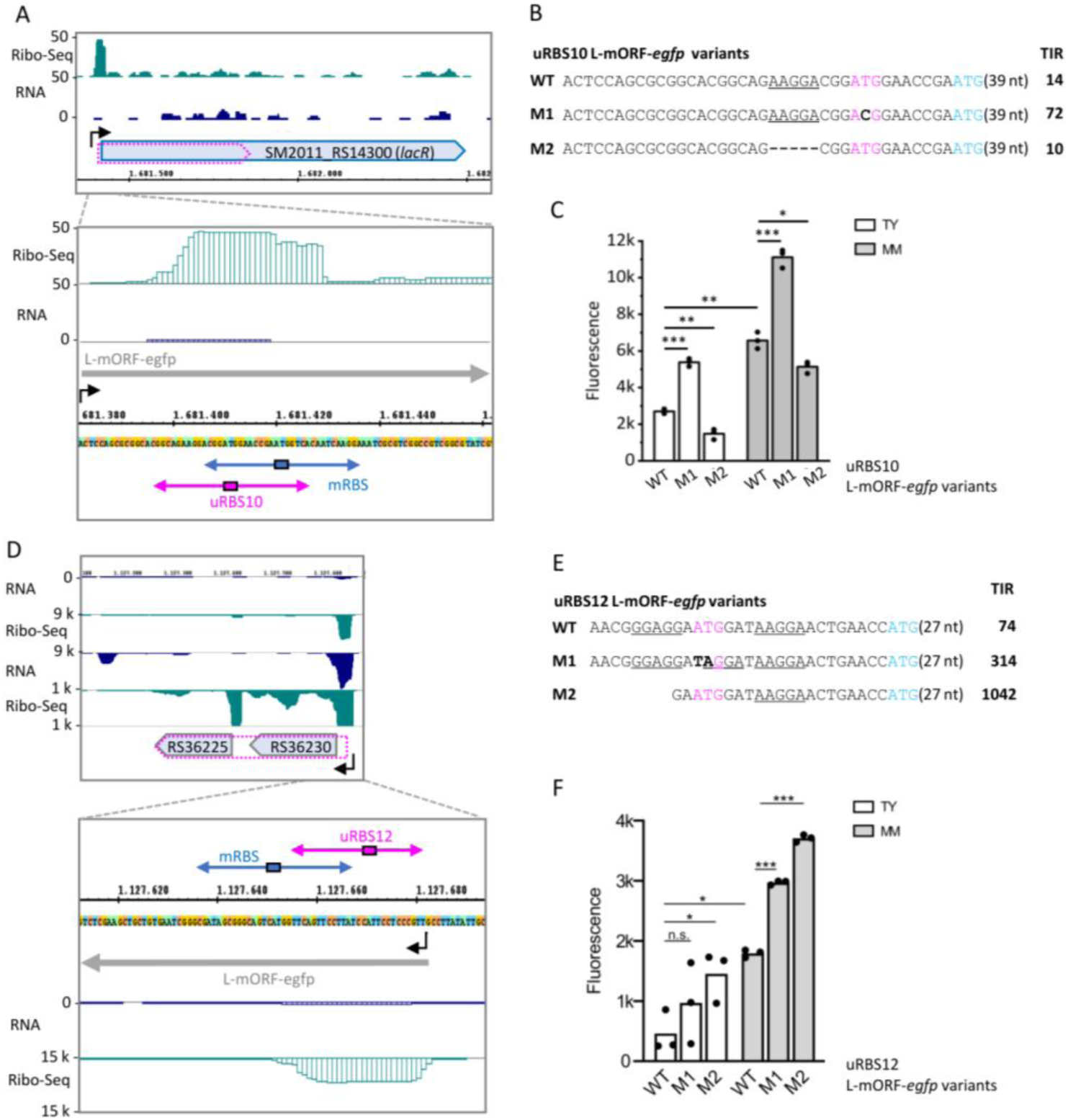
Ribosome occupancy at uRBS10 and uRBS12 interferes with mORF expression. **A)** and **D)** Integrated genome browser screenshots depicting reads from Ribo-seq and RNA-seq libraries. uORF10 (overlaps *lacR*) and uORF12 (overlaps the SM2011_RS36230-SM2011_RS36225 operon) are indicated by arrows with dashed pink lines. For other descriptions, see Fig. 3. The zoom-in panels show enrichment of Ribo-seq reads at uRBS10 and uRBS12. **B)** and **E)** Sequences fused to *egfp* in the indicated reporter constructs. Start codons are in pink for the uORF and cyan for the mORF. Shine-Dalgarno (SD) sequences that contain at least four nucleotides fully complementary to the anti-SD sequence of the *S. meliloti* 16S rRNA are underlined. Mutations are indicated in bold. Only the 5’-terminal regions of the cloned sequences are shown and their length is indicated (nt number in brackets). Predicted TIR values are given. **C)** and **F)** Fluorescence mediated by the indicated reporter constructs. All graphs show means and single data points of at least three independent experiments. Significance of difference determined by *t*-test: *** p ≤ 0.001, ** p ≤ 0.01, * p ≤ 0.05.

Further, we analyzed uRBS12, which did not promote uORF12-*egfp* translation (Fig. 2A and Fig. S2A above), but showed a very strong peak in Ribo-seq, indicating efficient occupancy by 70S ribosomes in MM conditions (Fig. 7D). The uORF12 fully overlaps with the mORF SM2011_RS36230, and corresponds to a larger isoform of the downstream gene SM2011_RS36225 (Fig. 7D). The distance between the uATG and the mATG is 16 nt and they do not share the same SD (see WT in Fig. 7E). Still, occupancy of uRBS12 by an initiating ribosome is expected to block the mRBS (Fig. 7D) and might serve in mORF regulation. For analysis of uORF12-eGFP translatability and uRBS12 role in mORF translation, the above strategy was used. The data show that when translated from a mutated RBS with favorable spacing between SD and start codon (Hyatt et al., 2010), the fusion protein uORF12-eGFP was detected (Fig. S7C and S7D). Furthermore, a start-to-stop mutation (M1) in the L-mORF-*egfp* fusion, which destroyed uORF12 (Fig. 7E), significantly increased mORF expression in MM (Fig. 7F). The M1 mutation created an artificial SD sequence AGGA, which is located 14 nt upstream of the mATG. However, since the mSD sequence AAGGA is located 8 nt upstream of the mATG (Fig. 7E), and this represents a better spacing between an SD and a start codon than a 14 nt-spacing (Hyatt et al., 2010), we suggest that the higher fluorescence of the M1 construct is due to the lack of ribosome binding to uRBS12 and thus unhindered binding to the mRBS. Moreover, a shortened mRNA leader lacking the uSD sequence (M2 mutation in Fig. 7E) also increased mORF expression (Fig. 7F), in line with our hypothesis that ribosome binding to uRBS12 occludes the mRBS and inhibits mORF translation.

### Provoked spatial conflict between uRBS and mRBS decreases mORF translation

To further test the hypothesis of uRBS-bound ribosomes that occlude mRBS accessibility, we decided to mutate the abovementioned L-mORF-*egfp* construct of uRBS1. It harbors the highly translated uORF1 (78 nt), which is translationally coupled to mORF1 by an ATGA overlap (see Fig. 5B above). We introduced deletions in uORF1, which do or do not provoke spatial conflict between uRBS1 and mRBS1, thereby preserving the ATGA overlap. When 39 nt of uORF1 are deleted (a Δ39 derivative), ribosome occupancy at the uRBS does not exclude occupancy at the mRBS (Fig. 8A). As expected, this Δ39 derivative mediated strong fluorescence, showing strong mORF1 translation (Fig. 8B). A deletion of additional 15 nt in uORF1 (a Δ54 derivative, Fig. 8A), leading to a spatial conflict between uRBS1 and mRBS1, resulted in significantly reduced fluorescence (compare Δ39 to Δ54 in Fig. 8B). This result is in line with a mutually exclusive ribosome occupancy at uRBS1 or mRBS1 in the Δ54 construct.

**Figure 8.**
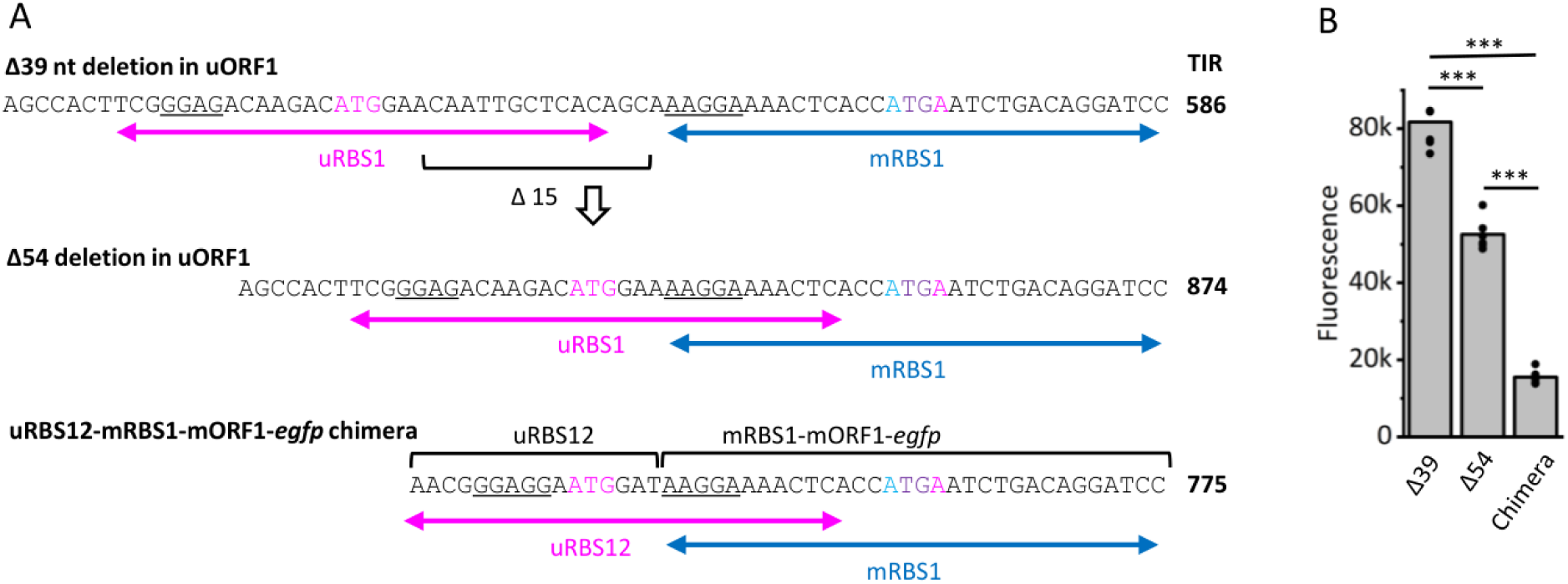
Provoked spatial conflict between uRBS and mRBS inhibits mRBS translation. **A)** Sequences fused to *egfp* in the indicated reporter constructs. The wild type sequence of L-mORF-*egfp*, which contains uRBS1, uORF1, and mORF1, is shown in Fig. 5B. Start codons are in pink for the uORF and cyan for the mORF. The start-stop overlap of uORF and mORF in ATGA is indicated in purple. Shine-Dalgarno (SD) sequences that contain at least four nucleotides fully complementary to the anti-SD sequence of the *S. meliloti* 16S rRNA are underlined. Horizontal double-headed arrows indicate the regions expected to be occupied by initiating ribosomes bound to the uRBS (pink) and mRBS (blue). Predicted TIR values are given. **B)** Fluorescence mediated by the reporter constructs, which are shown in A). Cultures grown in MM were used. The graph shows means and single data points of six independent experiments. Significance of difference determined by *t*-test: *** p ≤ 0.001, ** p ≤ 0.01, * p ≤ 0.05.

Additionally, we placed a 16 nt sequence harboring uRBS12 directly upstream of mRBS1, obtaining an uRBS12-mORF1-*egfp* chimera (Fig. 8A). This construct harbors a short chimeric uORF overlapping mORF1 by ATGA, and provokes a spatial conflict for ribosome occupancy on uRBS12 or mRBS1 (Fig. 8A). The significantly lower fluorescence of the chimeric construct in comparison to the Δ54 and Δ39 constructs demonstrates the strong inhibiting effect of uRBS12 (Fig. 8B).

### Re-screeining of uRBS predictions suggests additional uRBSs acting by direct mRBS occlusion

The above results suggest that closely spaced uRBS and mRBS could be used for regulatory purposes in bacteria. To address the question of how spread are closely spaced uRBS and mRBS in *S. meliloti*, Table S1 and Table S2 were re-screened to identify uRBS and mRBS pairs with start codon distances between 3 and 25 nt. We found 335 instances (Table S8) and have chosen three of them for experimental analysis. The three candidates have Ribo-seq profiles suggesting ribosome occupancy at the uRBS rather than at the mRBS (Fig. 9A to 9C). Their mORFs are first genes in operons and thus well suited for analysis by cloning uORF-*egfp* and L-mORF-*egfp* constructs, and by introducing start-to-stop mutations (designated M1; see Fig. 9D), which destroy the uORFs in the mRNA leaders of the latter.

**Figure 9.**
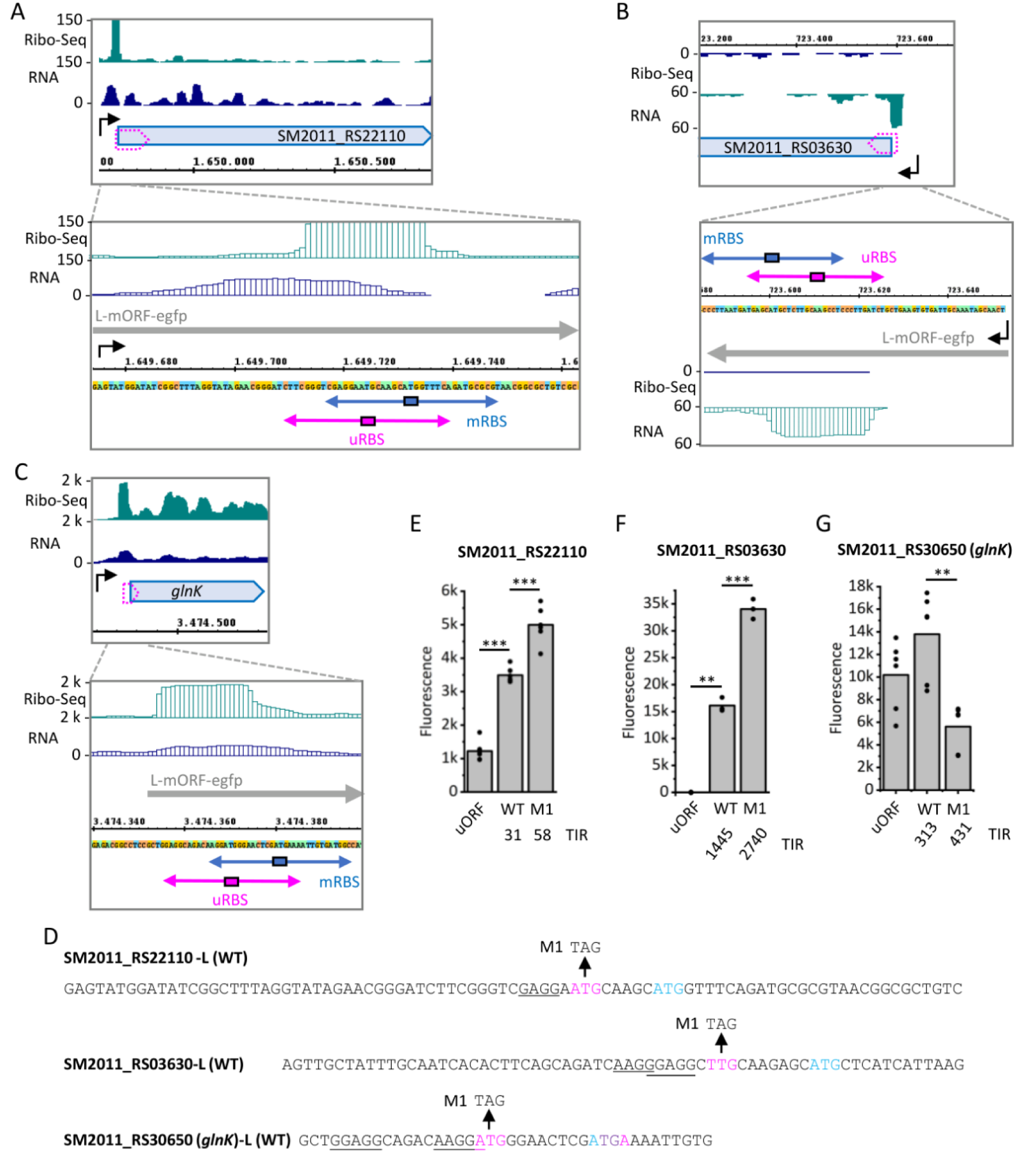
Re-screening of uRBS prediction reveals additional regulatory uRBSs that are closely spaced to the mRBSs. **A)** to **C)** Integrated genome browser screenshots depicting Ribo-seq and RNA-seq reads for the indicated genes. For other descriptions, see Fig. 3. **D)** Sequences fused to *egfp* in the used L-mORF-*egfp* constructs. The M1 mutation is indicated. Identical transcript starts were used for the uORF-*egfp* fusions. For other descriptions see Fig. 8. **E)** to **G)** Fluorescence mediated by the uORF-*egfp* fusions (uORF), and the WT and M1 variants of the L-mORF-*egfp* fusion shown in panel D. The predicted TIR values for mORF translation are given below the panels. MM cultures were used. All graphs show means and single data points of at least three independent experiments. Significance of difference determined by *t*-test: *** p ≤ 0.001, ** p ≤ 0.01, * p ≤ 0.05.

The gene SM2011_RS22110 (ABC transporter ATP-binding protein) has an mRNA leader of 57-nt harboring an uRBS. The uORF, which overlaps the mORF by 37 nt, shows relatively weak translation (Fig. 9E). The distance between the uATG and the mATG is 5 nt (Fig. 9D). The M1 mutation, which did not create an additional SD, significantly increased mORF translation (Fig. 9E). Similarly, the 51-nt mRNA leader of SM2011_RS03630 (alpha/beta fold hydrolase) harbors an uRBS of a putative uORF that overlaps the mORF by 23 nt. The distance between the uTTG and the mATG is 7 nt (Fig. 9D). No translation was detected for the uORF-*egfp*, and the M1 mutation increased mORF translation (Fig. 9F), without creating an additional SD (Fig. 9D). Both examples support the idea of closely spaced uRBS-mRBS pairs, in which the uRBS does not promote strong uORF translation, but efficiently binds ribosomes and attenuates the mORF expression.

Our attention was also drawn by *glnK* (SM2011_RS30650; P-II family nitrogen regulator), which has an ATGA-overlap with a 4-aa uORF and could be a subject of regulation by translational coupling. The uATG is separated from the mATG by 8 nt, and both start codons have strong SD sequences with suitable spacing (Fig. 9D). In this case, the uORF was well translated, and that the M1 mutation decreased the mORF translation (Fig. 9G), in line with the proposed translational coupling.

Thus, our re-screening of uRBS predictions, followed by experimental analysis, identified three additional uRBSs (uORFs), which affect mORF expression.

## DISCUSSION

Since the inventory of Ribo-seq (Ingolia et al., 2009), uORFs were found to be prevalent in mRNA leaders of eukaryotic genes and to exhibit a variety of regulatory functions, which are mechanistically mostly based on capturing the scanning ribosomal subunit and thus attenuating translation of the downstream mORF (Dever et al., 2023; Zhang et al., 2019). For many model bacteria, genome-wide identification of uORFs is still lacking. In this study, we used the alphaproteobacterium *S. meliloti* 2011 to predict uRBSs consisting of an uSD and a start codon, and detected uRBS candidates in the 100 nt-regions upstream of more than one thousand annotated genes. Among them are potential alternative or major TISs, as shown here in two examples and proposed for additional 23 genes (Fig. 3, Table S6). Thus, our study can contribute to improve the annotation of *S. meliloti* 2011, for which in different annotation releases, often different start codons were assigned to coding sequences (Hadjeras et al., 2023a). However, the majority of the uRBS candidates define putative uORFs. Our experimental data suggests that many of the uORFs could be translated and thus could impact expression of the downstream genes, or could encode functional small proteins. On the other hand, the detected uRBS candidates can be used by the evolution to test emerging regulatory uORFs, small genes or protein isoforms (Weaver et al., 2019; Zhong et al., 2023). Additionally, uRBSs harboring strong SD sequences could serve as binding sites for 30S ribosomal subunits for regulatory purposes (Jagodnik et al., 2025; Scheuer et al., 2022).

We note that a number of the predicted RBSs with a large distance to the mORF start probably are not uRBSs, because they are likely located upstream of TSSs. TSSs were not considered by OSTIR, but it is known that the majority of the mRNA leaders in *S. meliloti* are shorter than 100 nt (Schlüter et al., 2013). On the other hand, our prediction likely missed potential uRBSs and uORFs for several reasons. First, we considered only one uORF per mORF and did not account for mRNA leaders longer than 100 nucleotides (Bae and Crawford, 1990; Kothe et al., 2025). Second, our prediction of uRBSs was based on the presence of uSD sequences. However, even in bacteria that typically use the Shine-Dalgarno sequence for efficient translation initiation, many genes lack a recognizable SD sequence (Omotajo et al., 2015). Together with the results presented here, this suggests that bacterial uORFs are more prevalent than previously expected.

The experimental analysis of 16 uORF-defining uRBSs revealed potential roles for twelve of them (nine listed in Table 1 and three additional ones chosen after re-screening and shown in Fig. 9). Despite the limitations posed by our experimental approach (using reporter fusions on plasmids and under the control of a strong promoter; using artificial transcription starts in some of the constructs; lacking data on mRNA and protein stability differences between reporter constructs), we could assign functions to several new uRBSs and uORFs. The most plausible examples are the four uORFs that overlap the associated mORFs by few nucleotides and, according to our data (Fig. 5), act by translational coupling. Among others, our results suggest that in *S. meliloti*, a T1SS operon is translationally coupled to an uORF (Fig. 5A and 5B), which is regulated in response to the growth conditions (Fig. 2A and Fig. S2A). In rhizobia, T1SS secretes a nodulation factor, NodO, that shows homology to hemolysin (Scheu et al., 1992). Further, the chemotaxis gene *mcpW* (preceded by uORF11; Zatakia et al., 2018) and the P-II gene *glnK* (SM2011_RS30650; Fig. 9) are probably also translationally coupled to uORFs, since they have ATGA (start/stop) overlap (Brown and Wade, 2025). The 115 detected ATGA overlaps (Table S7) suggest that translational coupling is widespread in *S. meliloti*. As mentioned in the introduction, several mechanisms were proposed to mediate translational coupling. The 13 nt overlap between the uRBS9-defined uORF and the associated mORF suggests that their coupling is mediated by *de novo* binding of a 30S ribosomal subunit to the mRBS (Brown and Wade, 2025).

Among the analyzed uRBSs and uORFs are such having very high or very low predicted TIR and having ATG, GTG or TTG as start codons. According to the presented data, representatives with different TIR and different start codons have effects on the mORF expression (Table 1). Prominent examples are two positively acting uORFs uncovered in this work, one of them defined by the above mentioned uRBS1 (with high TIR and ATG start codon, translationally coupled to the T1SS operon), and the second one corresponding to uRBS8 (which has a very low TIR, defining the short GTG-uORF upstream of *hfq*). Although essentially no translation of this GTG-uORF was detected, a point mutation inactivating its start codon had a strong negative effect on the *hfq-egfp* expression (Fig. 6). A possible explanation of this result is that a 70S ribosome bound to the GTG-uORF masks an endonuclease cleavage site or a *rut* site utilized by the transcription termination factor Rho (Ben-Zvi et al., 2019; Kothe et al., 2025). Future analyses are needed to reveal the mechanisms of this uORF-based regulation of *hfq* encoding a major RNA chaperone (Santiago-Frangos and Woodson, 2018; Sobrero and Valverde, 2011).

Strikingly, our analysis revealed four uORFs with large overlaps to their mORFs, which, according to Ribo-seq, bind 70S ribosomes at their start codons, but are weakly or not translated (Fig. 7, Fig. 9 except *glnK*). They are characterized by uRBSs that are closely spaced to the mRBSs, which results in mutually exclusive occupancy by initiating ribosomes, and seems to attenuate the mORF expression. The here analyzed examples are characterized by a suboptimal (shorter than 4 nt) spacing between a potential uSD and the uORF start codon (Fig. 7, Fig. 9 except *glnK*). A possible interpretation of our data is that these uRBSs are occupied by ribosomes that inefficiently transit to elongation (we propose the name “sitting turtle” ribosomes). We are tempted to speculate that splitting the “sitting turtle” ribosomes under specific, yet unknown conditions, could increase the mORF expression.

According to the presented data, uRBS12 binds ribosomes and acts as a silencing element that suppresses expression of the first gene of a bicistronic operon of small genes with unknown function. The uORF12 corresponds to a larger protein isoform of the second gene in the operon (Fig. 7D). This is reminiscent of the regulation of toxin-antitoxin systems acting in phage defense (Zhong et al., 2023). Activating or splitting the ribosome at uRBS12 could change the expression of the whole operon. By contrast, uORF10 overlaps one-third of the *lacR* gene (Fig. 7A). The uORF10 shows expression in minimal medium (Fig. S2A) and might encode a functional protein needed under these conditions, while uRBS10 possibly fine-tunes the translation of *lacR*. Considering the large proportion of predicted uORFs with a large overlap with the associated mORFs (Fig. 1), many of them having uRBSs closely spaced to the mRBSs (Table S8), more *S. meliloti* genes could be attenuated by the proposed competition between uRBS and mRBS for mutually exclusive ribosome binding.

In summary, we mapped a genome-wide set of uRBS candidates in *S. meliloti* 2011 and provide examples for translated uORFs, which modulate the expression of the downstream genes. We further identified ribosome-binding uRBSs, which mediate weak or no translation, are in spatial conflict with the downstream mRBSs and thus attenuate mORF translation. These findings suggest that ribosome binding at uRBSs represents a novel mechanism for gene regulation at the level of RNA.

## ACKNOWLEDGEMENTS

We thank Leonie Schwaab, Chayma Chaib, and Alexandra Podkolodnich (Justus Liebig University Giessen) for help in some experiments. We acknowledge provision of computing resources by the Bioinformatics Core Facility (BCF) at Justus Liebig University Giessen.

## AUTHOR CONTRIBUTIONS

EEH, JMH and JB initiated the project. JMH conducted bioinformatic predictions and analysis. TD, SN, YAR, TW, SBW, and EEH designed the experiments, analyzed the data and conducted data visualization. JMH and EEH mainly wrote the manuscript with input and feedback from the other authors. AG, JB, and EEH supervised the research and provided resources and funding. All authors approved the submitted version.

## List of Supplementary Tables and Supplementary Figures

**Table S1: 912 predicted uRBSs in *S. meliloti* 2011.**

**Table S2: 211 predicted uRBSs with strong SD.**

**Table S3: Tool versions**

**Table S4: Used plasmids**

**Table S5: Used oligonucleotides**

**Table S6: Ribo-seq-supported uRBSs**

**Table S7: ATGA overlaps between predicted uORFs and their mORFs**

**Table S8: Closely spaced uRBS and mRBS pairs**

**Figure S1. Scheme of the uORF extraction and filtering workflow.** uORFs were extracted from the *S. meliloti* 2011 genome, and their translation initiation rates (TIRs) were predicted using OSTIR. Afterwards, uORFs overlapping with upstream annotated coding sequences (CDSs) were removed. For the remaining uORFs, the difference between the uORF and CDS/mORF TIRs was calculated (ΔTIR). Additionally, a Shine-Dalgarno motif search was performed. Upstream Shine-Dalgarno sequences (uSDs) overlapping with CDS/mORF were also excluded. uORF candidate lists were created from all uORFs with either a higher TIR compared to their CDS/mORF (Table S1 with 912 candidates) or a strongly conserved Shine-Dalgarno sequence “AGGAG” or “GGAGG” (Table S2 with 211 candidates).

**Figure S2. Correlation between predicted translation initiation rate (TIR) and fluorescence of bacterial cultures grown in MM as a read-out of ORF translatability in *Sinorhizobium meliloti*. A)** Analysis of 15 predicted uRBSs that represent putative uORF translation starts. Top: Scheme of the translational uORF-*egfp* fusions used for fluorescence (F) measurement. uSD: upstream Shine-Dalgarno sequence (SD), a part of the predicted upstream RBS (uRBS). uATG: start codon of the uRBS. The translational fusions are under the control of a strong heterologous promoter P*_metA_*. Bottom: Mean F values of bacterial cultures in minimal medium (MM), plotted against the predicted translation initiation rate (TIR) for each uRBS (uORF). For uRBS numbering, see Table1. **B)** Analysis of the corresponding annotated mORFs. Top: Scheme of the used translational mORF-*egfp* fusions. mSD, mATG: SD and start codon of the annotated mORF. Bottom: Mean F values of bacterial cultures grown in MM, plotted against the predicted TIR of each mORF. The mean F values represent data from three independent biological experiments, each measured in triplicates. In case the mean F value of a reporter construct was not significantly different from that of the empty vector control, F was set to 1. Results of Pearson and Spearman correlation analyses are shown. Coefficient of determination R^2^ and Spearman‘s Rho coefficient are given, and corresponding p-values are indicated: *** p ≤ 0.001, ** p ≤ 0.01, * p ≤ 0.05. Pearson regression line is shown and the confidence interval is indicated with dashed lines. The uRBS/uORF/mORF numbering is given in Table 1.

**Figure S3. Integrated genome browser screenshots depicting reads from Ribo-seq and RNA-seq libraries for the SM2011_RS08555 region.** Two transcription start sites (TSSs) are depicted by bent arrows. One of them corresponds to a leaderless SM2011_RS08555 transcript. The second TSS, annotated in the GenBank 2014 annotation, is located approximately 140 nt upstream and corresponds to a longer transcript with an mRNA leader harboring uRBS6. In the Ribo-seq library, read enrichment was detected upstream of uRBS6, suggesting the existence of a yet unanalyzed uORF in the mRNA leader, marked in gray. The region cloned in the L-mORF-*egfp* construct, which harbors uRBS6, is indicated by a horizontal gray arrow.

**Figure S4. Integrated genome browser screenshots depicting reads from Ribo-seq and RNA-seq libraries for uRBS9 (A), uRBS14 (B), and uRBS16 (C), and their associated mORFs.** The uORFs are indicated by pink arrows overlapping the blue mORF arrows, in which the RefSeq annotation numbers are given.

**Figure S5. Fluorescence of uRBS3, uRBS4, and uRBS13 L-mORF-*egfp* fusions and corresponding integrated genome browser screenshots depicting reads from Ribo-seq and RNA-seq libraries.** The Ribo-seq data refer to *S. meliloti* grown in MM (Hadjeras et al., 2023a). For mORF3, two alternative GTG start codons are indicated; the upstream one corresponds to a GenBank 2014 annotation. For legend description, see Fig. S4. WT: wild type construct. M1: derivative harboring a start-to-stop mutation destroying the uORF in the mRNA leader. M2: derivative with mutation in the uSD. The predicted TIR of the mORF in each construct is given at the bottom of the graphs. All graphs show means and single data points of at least three independent experiments. Significance of difference determined by *t*-test: *** p ≤ 0.001, ** p ≤ 0.01, * p ≤ 0.05.

**Figure S6. Analysis of uRBS2 and uRBS15. A)** and **C)** Integrated genome browser screenshots depicting reads from Ribo-seq and RNA-seq libraries for the indicated uRBSs and their associated mORFs. **B)** and **D)** Fluorescence mediated by the indicated constructs. uRBS15a corresponds to the predicted uRBS15 in Table 1. Here, an additional uRBS15b candidate was tested. All graphs show means and single data points of at least three independent experiments.

**Figure S7. uORF10 and uORF12 are weakly or not translated. A)** and **C)** show sequences cloned in translational uORF-*egfp* fusions. The uATG is in pink, SD sequences are underlined. Mutations (synonymous codons, non-synonymous codons, an insertions between the uSD and the uATG) are in bold. TIR values are given. **B)** and **D**) Fluorescence mediated by the corresponding constructs. Shown are means and single data points from at least three independent experiments. Growth conditions are indicated. Significance of difference determined by *t*-test: *** p ≤ 0.001, ** p ≤ 0.01, * p ≤ 0.05. **A)** and **B):** The uORF10-*egfp* variants 1, 2 and 3a differ by the number of uORF codons in the constructs. Although the predicted TIR of variant 1 (which is identical with the uORF10-*egfp* used in Fig. 2A and S2a) is lower than the TIR of variant 2, the latter showed no fluorescence. This could be explained by destabilization of the fusion protein by the 17 N-terminal uORF10 amino acids (aa). Variant 3a harbors four uORF10 aa fused to eGFP; it has low TIR and no fluorescence, and was used for further mutations. Variant 3b yielded fluorescence, which can be explained by the exchange of two rare codons synonymous, to non-rare ones. The non-synonymous codons exchange (variant 3c) did not mediate fluorescence, although the codons originate from the well translated uORF16-*egfp* (see Fig. 2A). Fluorescence was obtained when the sub-optimal 3-nt spacer between uSD and uATG was enlarged (variant 3d). **C)** and **D)** The non-synonymous codons used in variant 2 originating from the well translated uORF13 (see Fig. 2A and Fig. S2A). Variant 3: The spacer between uSD and uATG was enlarged using a spacer sequence from uORF13.

