## Supplemental Figures 1 to 7 for "Predicted Bacterial uRBSs Reveal Translational Coupling and Ribosome-Mediated RBS Occlusion as Gene-Controlling Mechanisms"

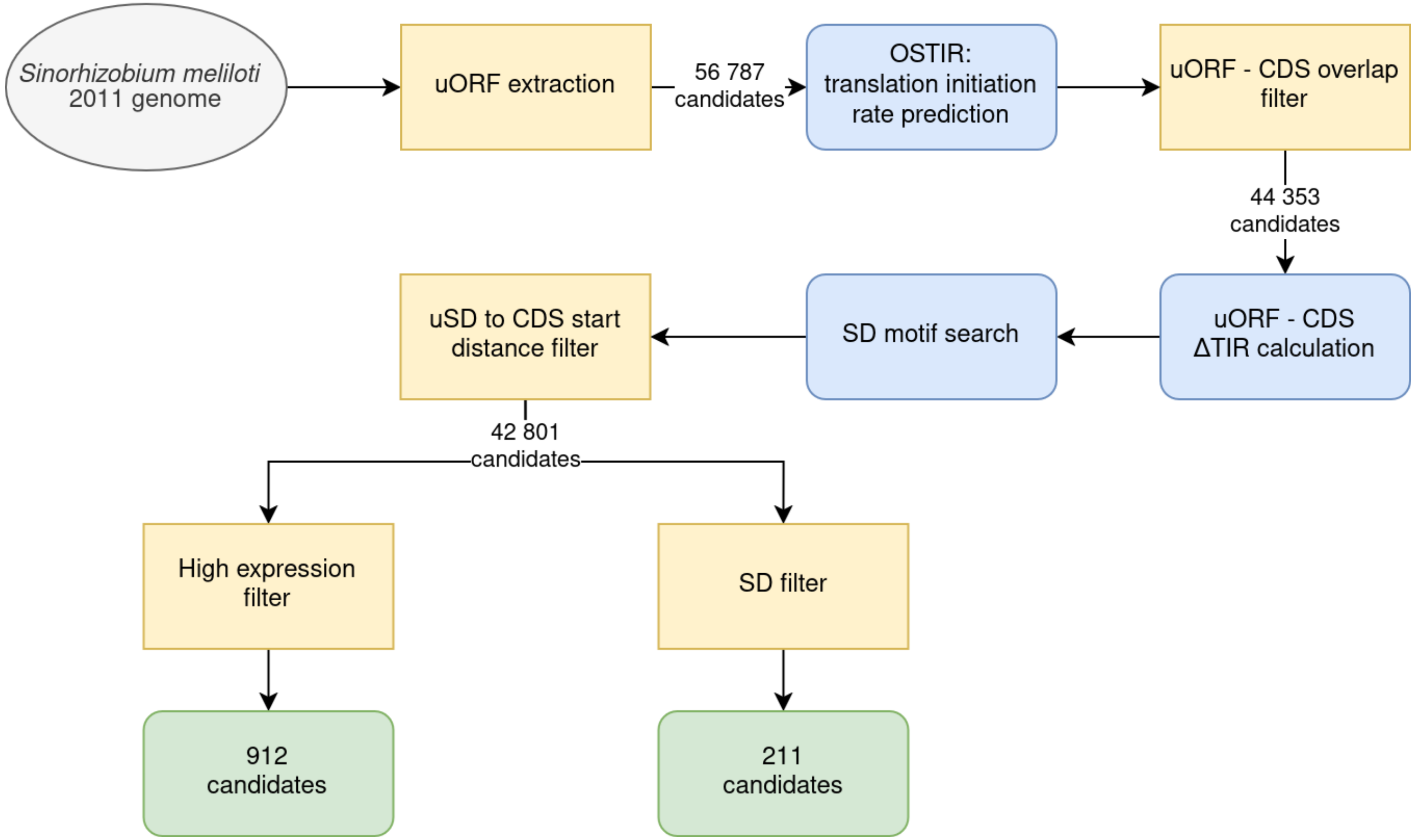


**Figure S1. Scheme of the uORF extraction and filtering workflow.** uORFs were extracted from the *S. meliloti* 2011 genome, and their translation initiation rates (TIRs) were predicted using OSTIR. Afterwards, uORFs overlapping with upstream annotated coding sequences (CDSs) were removed. For the remaining uORFs, the difference between the uORF and CDS/mORF TIRs was calculated (ΔTIR). Additionally, a Shine-Dalgarno motif search was performed. Upstream Shine-Dalgarno sequences (uSDs) overlapping with CDS/mORF were also excluded. uORF candidate lists were created from all uORFs with either a higher TIR compared to their CDS/mORF (Table S1 with 912 candidates) or a strongly conserved Shine-Dalgarno sequence "AGGAG" or "GGAGG" (Table S2 with 211 candidates).

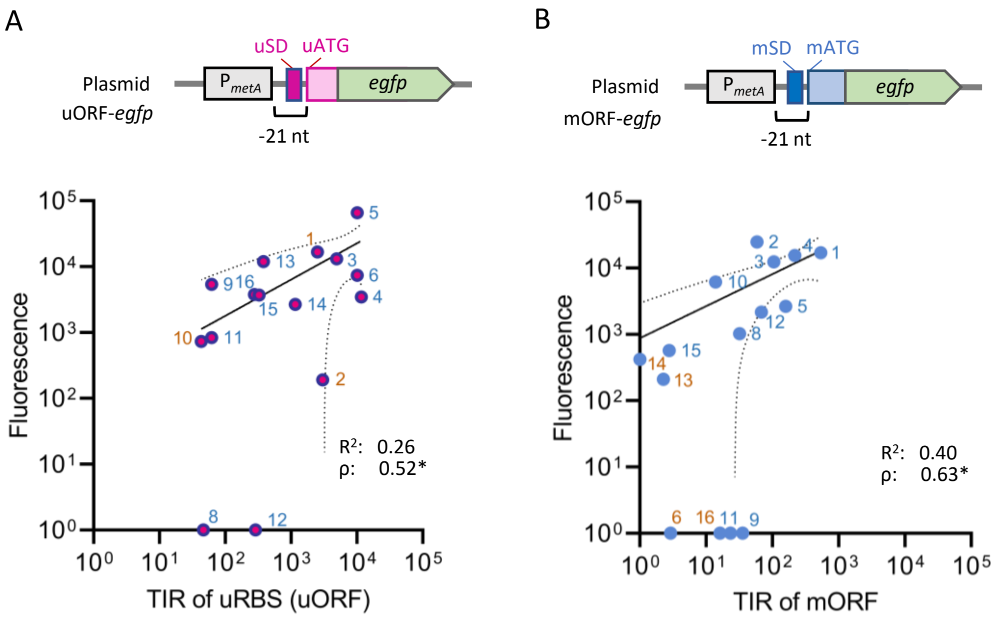


**Figure S2. Correlation between predicted translation initiation rate (TIR) and fluorescence of bacterial cultures grown in MM as a read-out of ORF translatability in *Sinorhizobium meliloti*.** **A)** Analysis of 15 predicted uRBSs that represent putative uORF translation starts. Top: Scheme of the translational uORF-*egfp* fusions used for fluorescence (F) measurement. uSD: upstream Shine-Dalgarno sequence (SD), a part of the predicted upstream RBS (uRBS). uATG: start codon of the uRBS. The translational fusions are under the control of a strong heterologous promoter $P_{metA}$. Bottom: Mean F values of bacterial cultures in minimal medium (MM), plotted against the predicted translation initiation rate (TIR) for each uRBS (uORF). For uRBS numbering, see Table1. **B)** Analysis of the corresponding annotated mORFs. Top: Scheme of the used translational mORF-*egfp* fusions. mSD, mATG: SD and start codon of the annotated mORF. Bottom: Mean F values of bacterial cultures grown in MM, plotted against the predicted TIR of each mORF. The mean F values represent data from three independent biological experiments, each measured in triplicates. In case the mean F value of a reporter construct was not significantly different from that of the empty vector control, F was set to 1. Results of Pearson and Spearman correlation analyses are shown. Coefficient of determination $R^2$ and Spearman's Rho coefficient are given, and corresponding p-values are indicated: *** $p \leq 0.001$, ** $p \leq 0.01$, * $p \leq 0.05$. Pearson regression line is shown and the confidence interval is indicated with dashed lines. The uRBS/uORF/mORF numbering is given in Table 1.

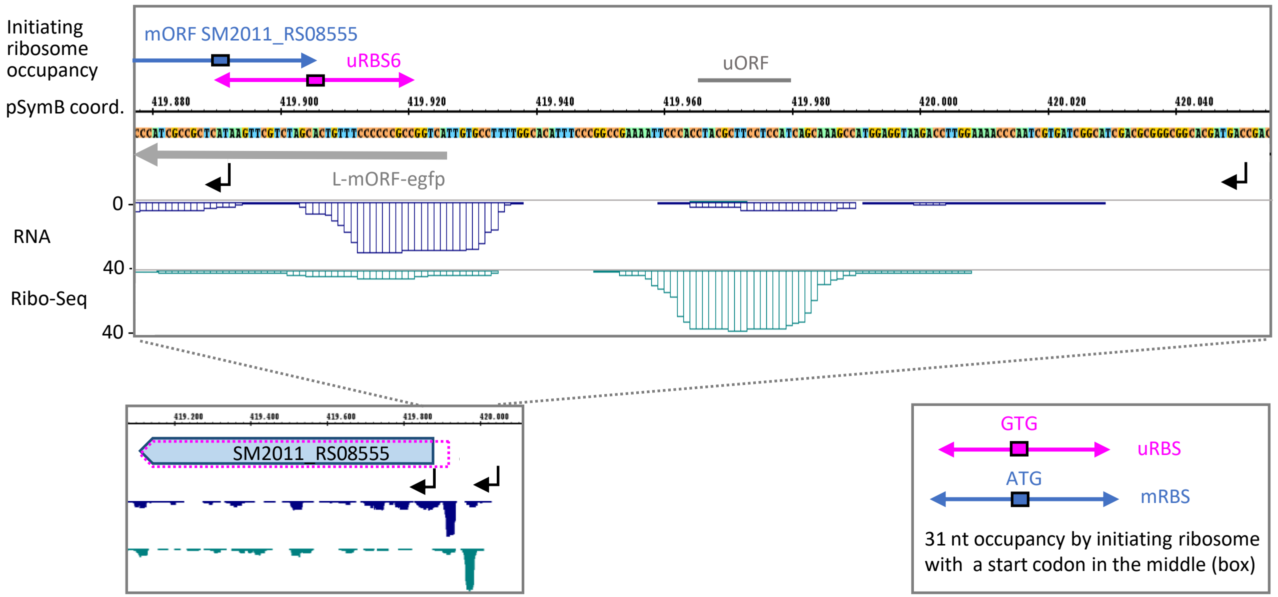


**Figure S3. Integrated genome browser screenshots depicting reads from Ribo-seq and RNA-seq libraries for the SM2011_RS08555 region.** Two transcription start sites (TSSs) are depicted by bent arrows. One of them corresponds to a leaderless SM2011_RS08555 transcript. The second TSS, annotated in the GenBank 2014 annotation, is located approximately 140 nt upstream and corresponds to a longer transcript with an mRNA leader harboring uRBS6. In the Ribo-seq library, read enrichment was detected upstream of uRBS6, suggesting the existence of a yet unanalyzed uORF in the mRNA leader, marked in gray. The region cloned in the L-mORF-*egfp* construct, which harbors uRBS6, is indicated by a horizontal gray arrow.

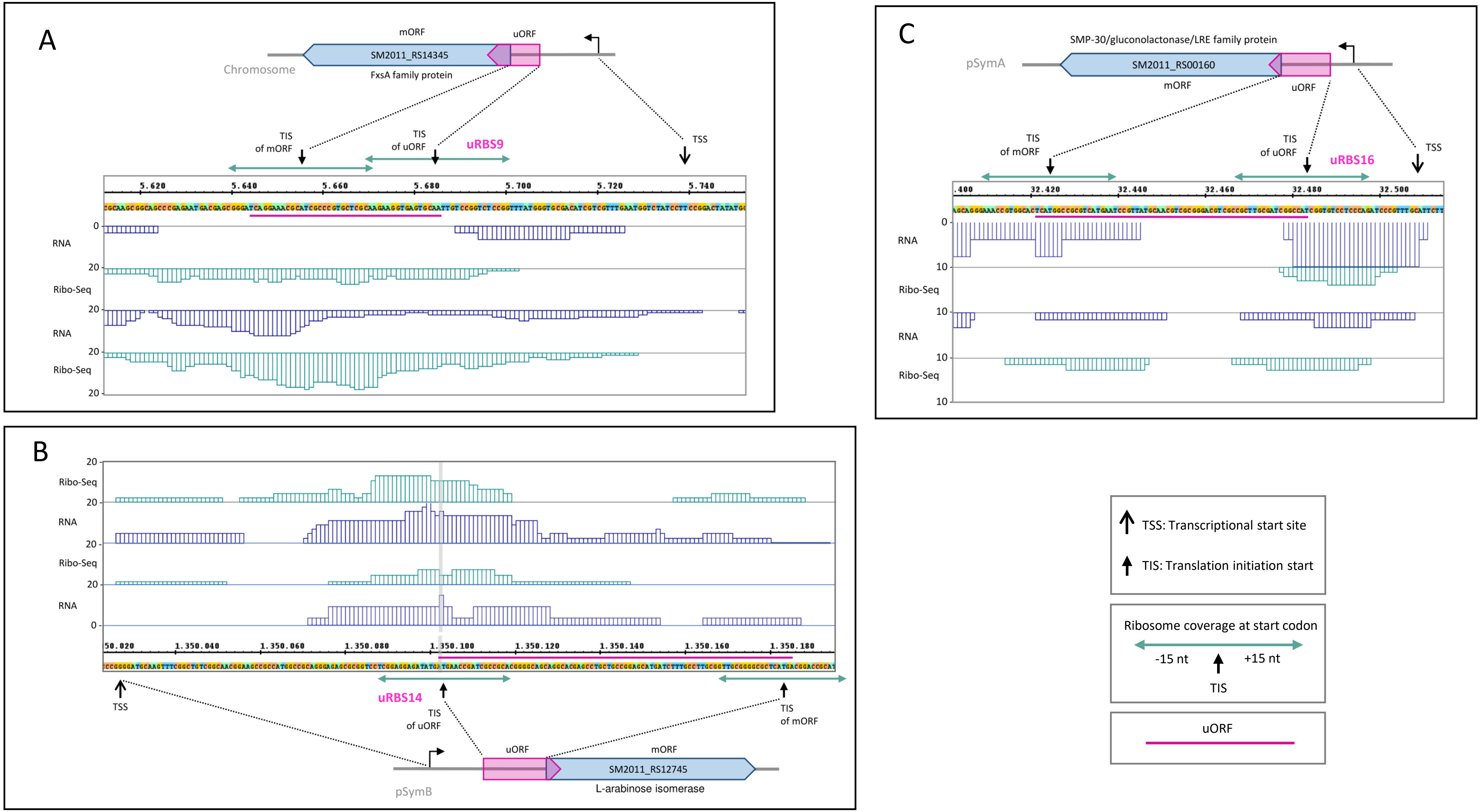


**Figure S4. Integrated genome browser screenshots depicting reads from Ribo-seq and RNA-seq libraries for uRBS9 (A), uRBS14 (B), and uRBS16 (C), and their associated mORFs.** The uORFs are indicated by pink arrows overlapping the blue mORF arrows, in which the RefSeq annotation numbers are given.

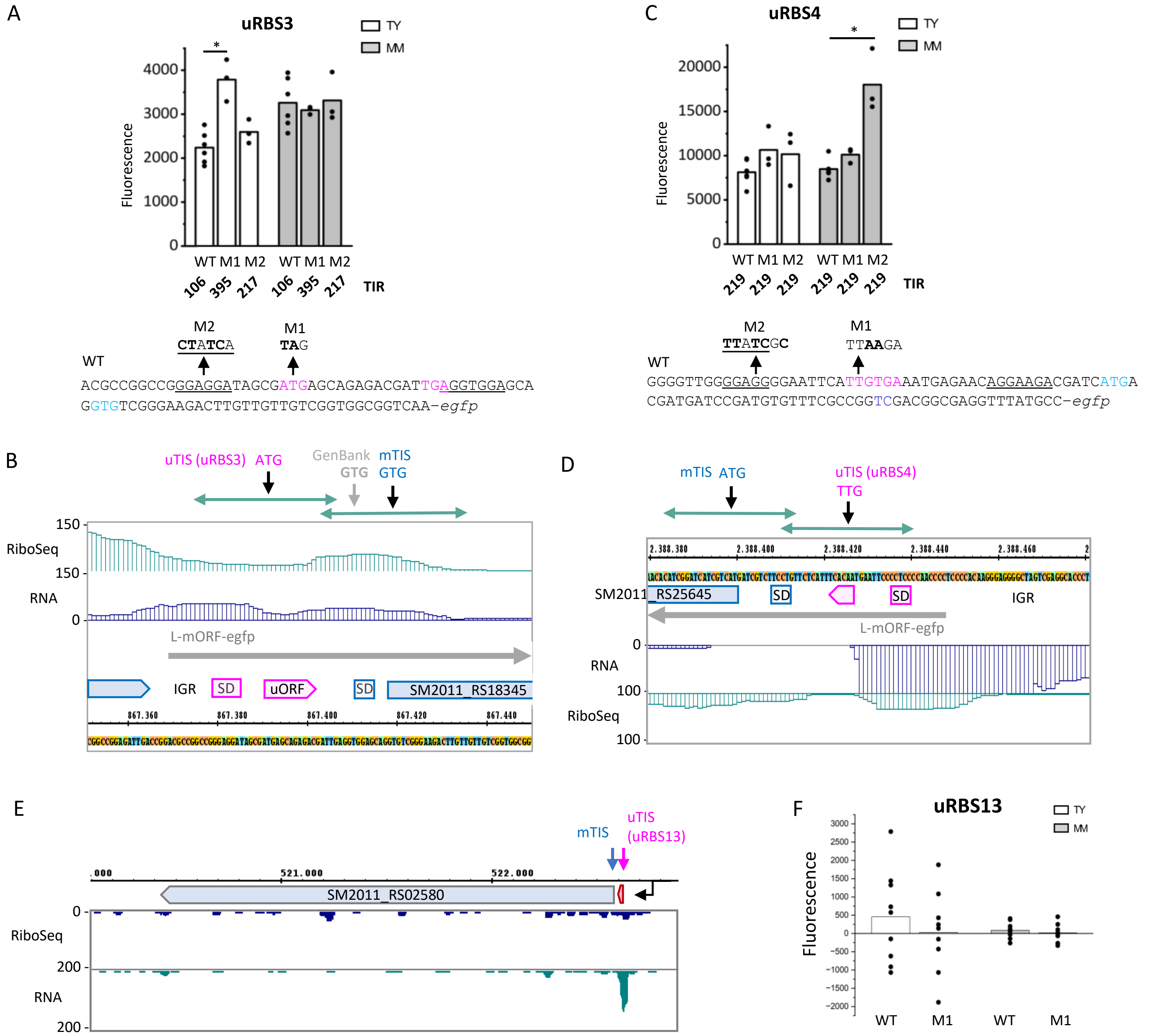


**Figure S5. Fluorescence of uRBS3, uRBS4, and uRBS13 L-mORF-*egfp* fusions and corresponding integrated genome browser screenshots depicting reads from Ribo-seq and RNA-seq libraries.** The Ribo-seq data refer to *S. meliloti* grown in MM (Hadjeras et al., 2023a). For mORF3, two alternative GTG start codons are indicated; the upstream one corresponds to a GenBank 2014 annotation. For legend description, see Fig. S4. WT: wild type construct. M1: derivative harboring a start-to-stop mutation destroying the uORF in the mRNA leader. M2: derivative with mutation in the uSD. The predicted TIR of the mORF in each construct is given at the bottom of the graphs. All graphs show means and single data points of at least three independent experiments. Significance of difference determined by *t*-test: *** p ≤ 0.001, ** p ≤ 0.01, * p ≤ 0.05.

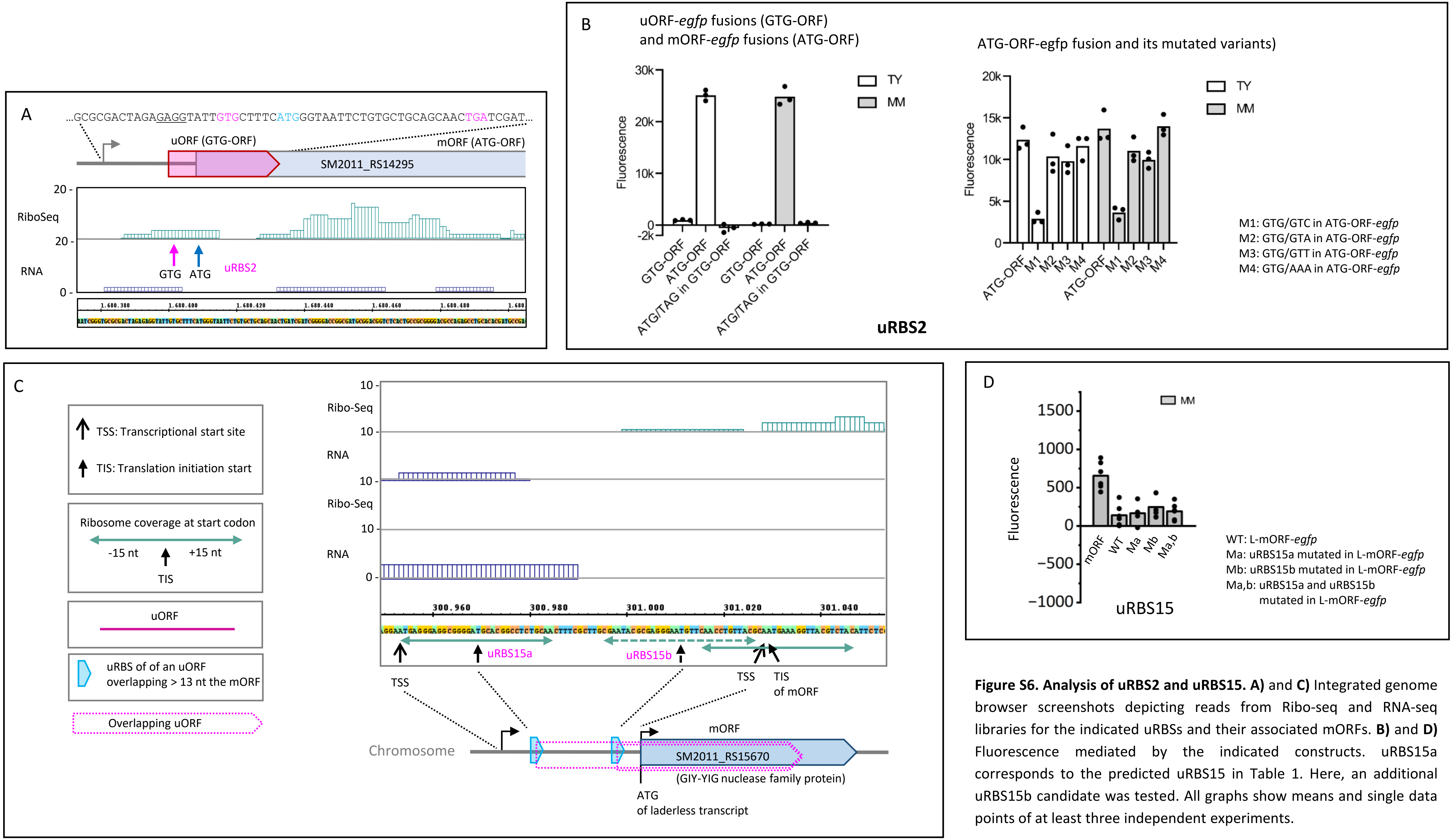


**Figure S6. Analysis of uRBS2 and uRBS15. A)** and **C)** Integrated genome browser screenshots depicting reads from Ribo-seq and RNA-seq libraries for the indicated uRBSs and their associated mORFs. **B)** and **D)** Fluorescence mediated by the indicated constructs. uRBS15a corresponds to the predicted uRBS15 in Table 1. Here, an additional uRBS15b candidate was tested. All graphs show means and single data points of at least three independent experiments.

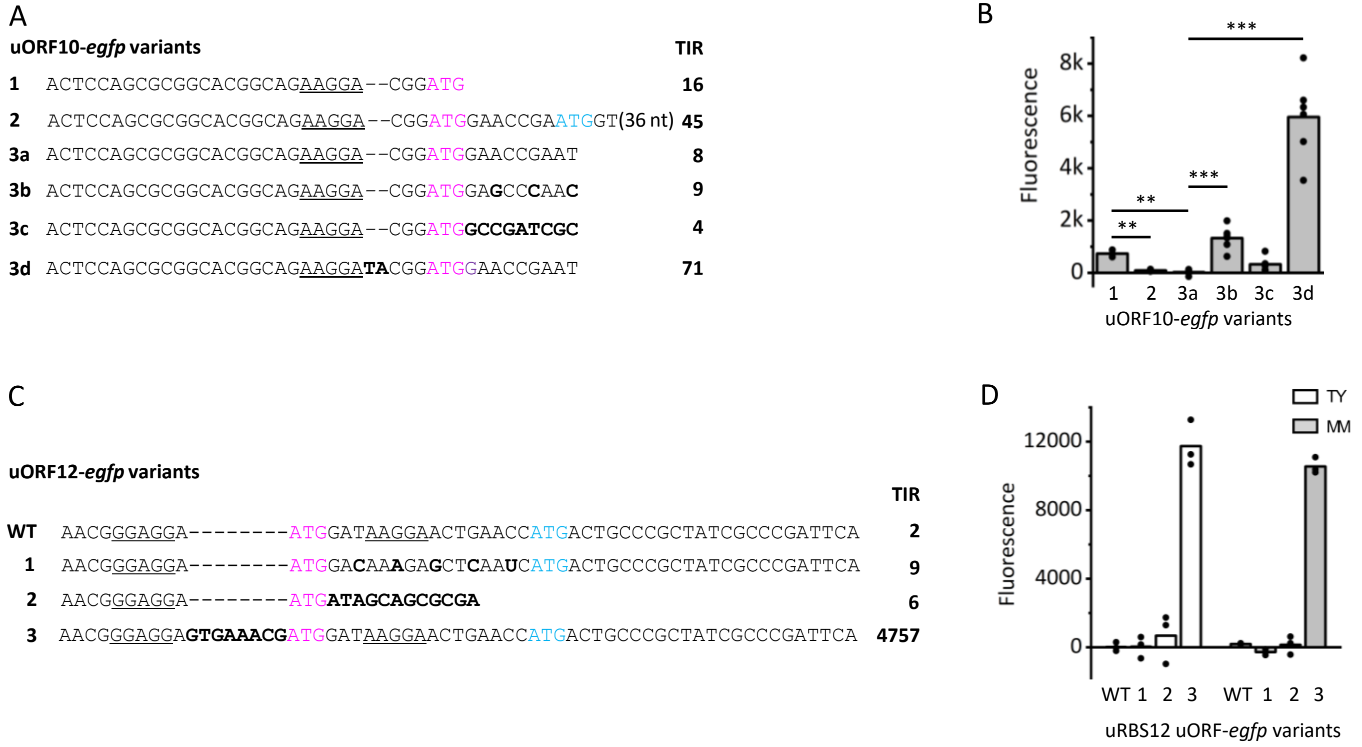


**Figure S7. uORF10 and uORF12 are weakly or not translated. A)** and **C)** show sequences cloned in translational uORF-*egfp* fusions. The uATG is in pink, SD sequences are underlined. Mutations (synonymous codons, non-synonymous codons, an insertions between the uSD and the uATG) are in bold. TIR values are given. **B)** and **D)** Fluorescence mediated by the corresponding constructs. Shown are means and single data points from at least three independent experiments. Growth conditions are indicated. Significance of difference determined by *t*-test: *** p ≤ 0.001, ** p ≤ 0.01, * p ≤ 0.05.

**A)** and **B)**: The uORF10-*egfp* variants 1, 2 and 3a differ by the number of uORF codons in the constructs. Although the predicted TIR of variant 1 (which is identical with the uORF10-*egfp* used in Fig. 2A and S2a ) is lower than the TIR of variant 2, the latter showed no fluorescence. This could be explained by destabilization of the fusion protein by the 17 N-terminal uORF10 amino acids (aa). Variant 3a harbors four uORF10 aa fused to eGFP; it has low TIR and no fluorescence, and was used for further mutations. Variant 3b yielded fluorescence, which can be explained by the exchange of two rare codons synonymous, to non-rare ones. The non-synonymous codons exchange (variant 3c) did not mediate fluorescence, although the codons originate from the well translated uORF16-*egfp* (see Fig. 2A). Fluorescence was obtained when the sub-optimal 3-nt spacer between uSD and uATG was enlarged (variant 3d).

**C)** and **D)** The non-synonymous codons used in variant 2 originating from the well translated uORF13 (see Fig. 2A and Fig. S2A). Variant 3: The spacer between uSD and uATG was enlarged using a spacer sequence from uORF13.
